# Dispersal, gene flow and adaptive differentiation in hierarchically structured populations

**DOI:** 10.64898/2026.09.16.751951

**Authors:** Emilio Egal, Mathieu Buoro, Aurélie Manicki, Guillaume Evanno, Charles Perrier

**Affiliations:** DECOD, INRAE, Institut Agro, IFREMER, Rennes, France; Management of Diadromous Fish in their Environment, OFB, INRAE, Université de Pau et des Pays de l’Adour, Institut Agro, Saint-Pée-sur Nivelle, France; Université de Pau et des Pays de l’Adour, INRAE, ECOBIOP, Saint-Pée-sur-Nivelle, France; CBGP, INRAe, CIRAD, IRD, Montpellier SupAgro, Université de Montpellier, Montpellier, France

**Keywords:** Local adaptation, gene flow, dispersal, spatial scale, hierarchical structure, Salmo salar

## Abstract

The balance between local adaptation and gene flow is rarely quantified when populations are hierarchically differentiated across multiple spatial scales. We addressed this gap in Atlantic salmon (*Salmo salar*) at its southern range edge, using 1342 individuals from 21 rivers along the French Atlantic and Channel coasts with a 60k SNP array, and combining estimates of dispersal with genome-wide scans for adaptive differentiation across spatial scales. Population structure was hierarchical, resolving five genetic clusters with differentiation among groups five times higher than among populations within them. Assignment tests revealed a marked contrast between clusters, but unassigned individuals reached very high proportions within some clusters, indicating that gene flow has already blurred among-populations differences. Adaptive differentiation was detected among and within clusters, but the loci involved overlapped only partially across scales and among clusters, indicating that adaptation at each hierarchical level relies on distinct sets of variants. Dispersal was a poor guide to where this signal persisted: retaining putative dispersers reduced the candidate set more than threefold, showing that immigrant genotypes could dilute among-populations allele-frequency contrasts within a single generation. Yet differentiation among populations remained strong, as expected if dispersers reproduce poorly. Resolving the balance between local adaptation and gene flow therefore requires accounting for dispersal and adaptive differentiation at different spatial scales, with direct implications for conserving hierarchically structured populations at climatically vulnerable range edges.

## Introduction

Local adaptation — the higher inherited fitness of individuals in their native environment relative to individuals originating elsewhere (Kawecki & Ebert 2004) — can be influenced by gene flow as a double-edged force. Depending on its strength relative to divergent selection, gene flow can either dilute locally advantageous alleles until adaptive differentiation is erased, or replenish genetic variation and prevent the erosion of diversity (Blanquart *et al*. 2013; Lenormand 2002; Tigano & Friesen 2016; Yeaman & Otto 2011). Which outcome prevails depends on the migration–selection balance, and that balance cannot be read from dispersal rates alone: it is shaped by life history, landscape heterogeneity, and the spatial scale at which it is observed (Fronhofer *et al*. 2024; Richardson *et al*. 2014; Savolainen *et al*. 2007). At which spatial scale, and through which genomic mechanism, local adaptation withstands gene flow therefore remains an open question.

A first obstacle lies in the distinction between dispersal and gene flow (i.e. effective dispersal). A disperser only contributes to the evolution of the recipient gene pool if it successfully reproduces there; dispersal and gene flow can therefore diverge substantially depending on the relative reproductive success of immigrants and residents (Egal *et al*. 2026; Fraser *et al*. 2011; Reid *et al*. 2024). Dispersal does not translate necessarily into gene flow, and the gap between the two is expected to widen in hierarchically structured systems, where gene flow declines and environmental contrasts increase with geographic distance (Alther *et al*. 2021; Brauer *et al*. 2018; Mullen *et al*. 2010). Measuring dispersal and adaptive differentiation in the same populations is therefore a prerequisite for interpreting either.

A second obstacle lies in the genomic architecture of adaptation. Theory predicts that under migration–selection balance, the architecture of adaptive loci evolves toward concentration in a few tightly linked regions, because such architectures resist homogenisation by gene flow more effectively than a diffuse polygenic basis (Yeaman & Whitlock 2011). In several fishes, chromosomal rearrangements and regions of reduced recombination do protect adaptive variants from homogenisation (Barth *et al*. 2017; Lehnert *et al*. 2020; Wellband *et al*. 2019). However, concentrated architecture and polygenic variation are not competing hypotheses: they are two axes along which the genomic response to divergent selection can vary, and which jointly determine whether — and where in the genome — local adaptation survives ongoing gene flow (Bernatchez 2016; Pritchard & Di Rienzo 2010; Pritchard *et al*. 2018).

Atlantic Salmon (*Salmo salar* Linnaeus, 1758) offers a particularly suitable system for disentangling these two obstacles. Its anadromous life cycle theoretically allows substantial dispersal among demes at both short and long distance, but strong homing behaviour returns most individuals to their natal river, maintaining significant genetic differentiation among populations (Hasler & Scholz 1983). Across salmonids, residents commonly outperform immigrants in the recipient river (Egal *et al*. 2026; Mobley *et al*. 2019; Peterson *et al*. 2014); yet dispersal remains far from negligible, ranging from below 1% in some Pacific species to over 27.5% in one Spanish *S. salar* population (Consuegra *et al*. 2005; Jonsson *et al*. 2003; Quinn & Fresh 1984; Stabell 1984), which makes the gap between dispersal and gene flow a likely key mechanism maintaining population differentiation in this species (Fraser *et al*. 2011; Lamarins *et al*. 2024a). In addition, fine-scale selection signatures have accordingly been found, notably linked to genes involved in sexual maturation, energy homeostasis and immune defence (*vgll3*, *mc4r*, major histocompatibility complex; Lehnert *et al*. 2020; Pritchard *et al*. 2018), while regional-scale landscape analyses have connected adaptive divergence to temperature and geological gradients (Bourret *et al*. 2013). Whether the same genomic regions are identified across different spatial scales in a given system, however, remains largely unknown.

That caveat matters because population structure in this species is explicitly hierarchical: among tributaries within rivers (Dillane *et al*. 2008; Vähä *et al*. 2007), among neighbouring rivers (Fontaine *et al*. 1997), and among broader regional groups (Bourret *et al*. 2013; Dionne *et al*. 2008). In France, microsatellite analysis of 975 individuals from 34 rivers resolved five geographically coherent clusters, with coastal distance, geological substrate and river length contributing to differentiation, and with gene flow highest among geologically similar rivers (Perrier *et al*. 2011). The scales so delimited differ in both selective and demographic terms. Clusters are separated by broad climatic and geological gradients, whereas neighbouring rivers sharing a macroclimate may differ mainly in fine-scale hydrology while exchanging a substantial fraction of their spawners within a single generation (Fontaine *et al*. 2025; Jonsson *et al*. 2003). There is therefore no reason to expect the same genomic targets at both scales — the genetic basis of a trait can be parallel or non-parallel depending on the scale compared (Kess *et al*. 2024) — and a signal recovered at one scale cannot be assumed to reflect the same process as a signal recovered at another. This question is especially relevant at the species’ southern, warm range edge, where populations are expected to be particularly sensitive to climate change (Valiente *et al*. 2010). Gabián *et al*. 2022 showed that Iberian salmon differ adaptively from Scottish populations, with signals linked to development and cellular metabolism — but their comparison involved broad geographic groups and did not quantify contemporary dispersal within a hierarchical river network.

Here we used a 60K SNP array to genotype 21 Atlantic salmon populations along the French Atlantic and English Channel coasts, at the southern edge of the range. Our aim is to examine how dispersal, gene flow, hierarchical structure and local adaptation interact across spatial scales, and how the resulting adaptive signal is distributed along the genome. We first revisit the hierarchical organisation of these populations at the genome-wide level. We then identify contemporary dispersers using population assignment tests, and ask whether signatures of adaptive differentiation persist among and within clusters once these individuals are accounted for. Finally, we test whether putative adaptive differentiation is associated with river-scale environmental variation, while accounting for neutral population covariance.

## Materials and Methods

### Study site and population sampling

We sampled 1506 Atlantic salmon (*Salmo salar*) individuals from 21 sites along the Bay of Biscay and English Channel (Table 1). To ensure a robust population assignment, we collected both juveniles and adults. Juveniles were caught by electrofishing and adults by angling or using trapping stations (only for BID, BRE, NVL, SCO, SEL). Fin clips and scales were collected for each individual and preserved in 95% ethanol and in paper envelopes, respectively.

**Table 1.**
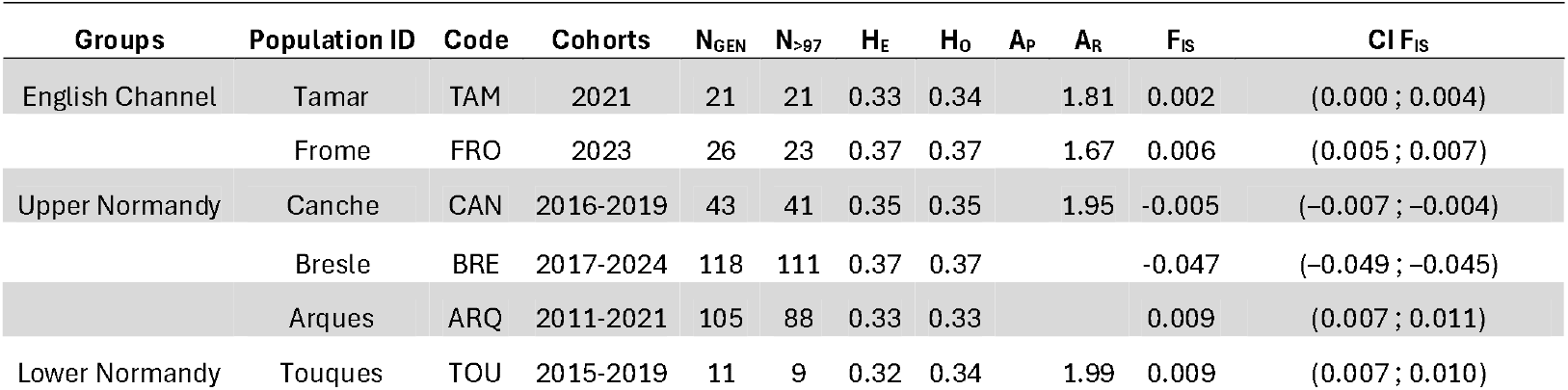
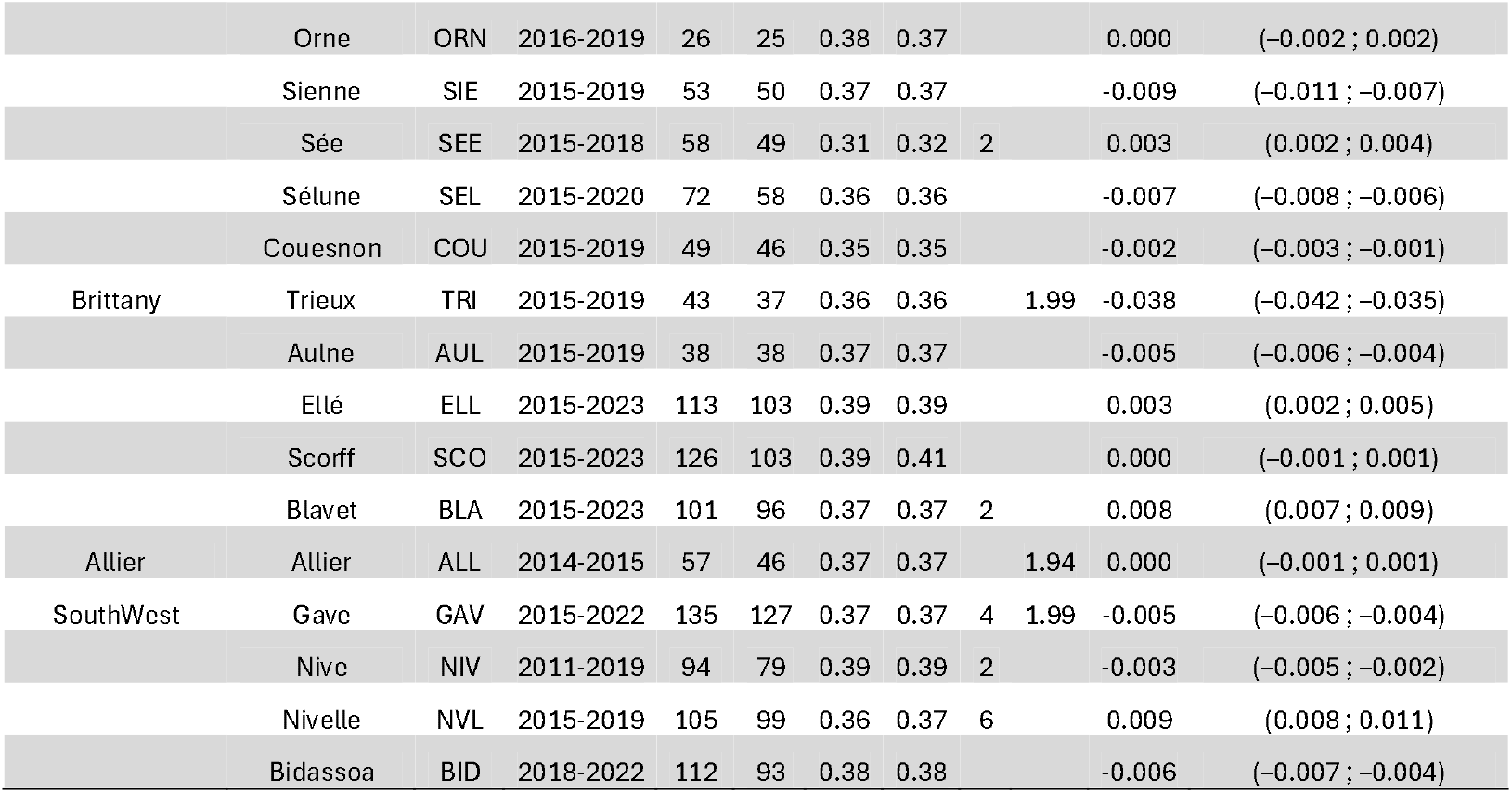
Sampling and genetic diversity summary for the 22 Atlantic salmon river populations analysed. Populations are ordered from north to south and assigned to regional groups (English Channel, Upper Normandy, Lower Normandy, Brittany, Allier, South-West). Columns are: Population ID and three-letter code; Cohorts, range of sampling years; NGEN, number of individuals genotyped; N_>97_, number retained after filtering on genotyping call rate (>97%); H_E_ and H_O_, expected and observed heterozygosity; A_P_, number of private alleles; A_R_, allelic richness by clusters; F_IS_, inbreeding coefficient, with its 95% confidence interval (CI F_IS_).

### Molecular analyses and genotyping

We extracted genomic DNA with the NucleoSpin 96 Tissue kit, quantified it with a Qubit 2.0 fluorometer and standardised its concentration to ≥23 ng/μl. We checked DNA integrity on agarose gel. Standardized DNA samples were genotyped at the Center for Integrative Genetics (CIGENE, Ås, Norway) on a custom 60K Atlantic salmon SNP array derived from a 220K Affymetrix Axiom platform (Barson *et al*. 2015).

We retained SNPs with call rates >0.97 and MAF >0.01 (n = 49,230) stored in our "PolyHighRes" dataset. We removed 47 mitochondrial and 21 unplaced markers. We retained individuals with call rates >97%, and we kept each unique multilocus genotype once, using mlg.filter in poppr (Kamvar *et al*. 2014) to identify related individuals and diss.dist to estimate genotyping error from technical replicates (Hamming distance). The final dataset comprised 1342 individuals (739 adults, 603 juveniles) and 49,162 SNPs, with a mean genotyping error rate of 0.2%. For the analysis of genetic structure only, SNPs were LD-pruned in PLINK v1.9 (-- indep-pairwise 50 5 0.5; (Chang *et al*. 2015), yielding 48,933 independent markers for structure analyses.

### Population structure, genetic diversity and effective population size

We ran a Principal Component Analysis (PCA) in pcadapt v4.4.0 (Privé *et al*. 2020; K = 4 retained from scree plots, Fig. S3), an individual-based clustering analysis in ADMIXTURE (Alexander *et al*. 2009; K = 1–10, selected by cross-validation error), and a Discriminant Analysis in Principal Components in adegenet (Jombart *et al*. 2010; BIC-based cluster number, 1,500 PCA axes, α-score optimisation retaining 13 PCs). We built a neighbour-joining tree of Nei’s genetic distances (Nei *et al*. 1983) with 1,000 bootstrap replicates (aboot, poppr). Pairwise F*_ST_* (Weir & Cockerham 1984) was estimated in hierfstat (Goudet 2005), with 95% confidence intervals from 1,000 bootstraps over loci. We tested the hierarchical partitioning of genetic variance with an AMOVA (poppr v2.9.8) across nested biogeographical groups, excluding Allier owing to its isolation and divergence, with significance assessed by 999 permutations (randtest, ade4; (Dray & Dufour 2007). Within-population genetic diversity was characterised in hierfstat from expected (H*_e_*) and observed (H*_o_*) heterozygosity, and from the inbreeding coefficient F*_IS_*, for which 95% confidence intervals were obtained from 1,000 bootstraps over loci. Private alleles were identified per population in poppr. We estimated contemporary effective population size (N*_e_*) with the LD-based method in CurrentNe2 (Santiago *et al*. 2025), which models subdivision and migration; for each population we compared inter-chromosomal panmictic estimates to a two-patch subdivision-with-migration model (CurrentNe2, ’-x’ option), retaining the subdivision-based N*_e_*, together with its associated migration rate m, when N*_e_* (structure) exceeded the inter-chromosomal panmictic estimate with non-overlapping 90% confidence intervals (Table S2).

### Identification of dispersers

We inferred the natal origin of adults using population assignment in assignPOP v1.3.0 (Chen et al. 2018), treating juveniles as baseline source populations. We ran analyses separately for four geographic groups — Southwest, Brittany, Lower Normandy and Upper Normandy — because dispersal is expected to decline with distance (Jonsson et al. 2003). We evaluated baseline performance with a Support Vector Machine classifier using all SNPs (no locus subsetting, to avoid marker high-grading bias; Anderson 2010), via Monte-Carlo cross-validation (60/75/90% training proportions, 30 replicates) and 5-fold cross-validation. For Southwest and Upper Normandy we additionally built refined baselines by removing juveniles consistently misassigned across k-fold replicates. Because the same cross-validation was used both to select and to evaluate these individuals, refined-baseline accuracy should be treated as an upper bound rather than an unbiased estimate. We considered adults with assignment probability q ≥ 0.70 as confidently assigned; individuals below this threshold were considered unassigned. We computed river-level dispersal rates as the proportion of adults confidently assigned to a different source population. We conducted a separate global (instead of regional) assignment analysis, using all juvenile populations as sources, plus adults from two rivers lacking juvenile samples (Allier, Touques), and including Tamar and Frome juveniles as sources (previously excluded from the regional analyses).

### Adaptive differentiation and cluster-dependent associations

Because river-network topology can inflate neutral F*_ST_* variance and produces high false-positive rates in standard outlier tests (Fourcade *et al*. 2013), we used methods that explicitly model the covariance structure of population allele frequencies, i.e. XtX statistic (Günther & Coop 2013) implemented in BayPass v3.1 (Gautier 2015). We removed putative dispersers (q ≥ 0.70) prior to all genome scans to avoid distorting local allele-frequency estimates. However, we also ran a complementary XtX scan retaining dispersers to assess their effect on outlier detection. Global genome scans used BayPass v3.1 (Gautier 2015) under the core model, computing the XtX statistic (Günther & Coop 2013) — an F*_ST_* corrected for the scaled covariance matrix Ω — on the unpruned, 49,162-SNP dataset. Each run comprised 25 pilot runs (1,000 iterations), a 100,000-iteration burn-in, and 2,500 sampling iterations (thinning = 40). Five independent runs were highly consistent (low Förstner–Moonen distance between Ω matrices; pairwise XtX correlations > 0.995; Table S4A-B), so we only reported the first run (seed 5100). We calibrated XtX values against a posterior-predictive null from 100,000 pseudo-observed SNPs (POD), with candidates defined as those exceeding the 99.5% quantile. We further applied a local-score approach (Fariello *et al*. 2017; threshold ξ = 2) to XtX p-values to delineate contiguous outlier-enriched genomic windows. We repeated the same procedure within each of the four geographic groups (within-cluster scans) using the local-only dataset; because Ω differs among these narrower analyses, XtX magnitudes are not directly comparable across scans, and comparisons were restricted to outlier identity and genomic position, visualised with an UpSet plot (UpSetR; Conway *et al*. 2017). To detect cluster-specific signatures, we computed C2 statistics contrasting each cluster against all others pooled (Southwest, Brittany, Lower Normandy, Upper Normandy, Allier), again consistent across five runs (correlations > 0.995; Table S4C) and visualised with an UpSet plot. To attribute global-scale outlier signals to specific populations, we ranked populations by the absolute standardised posterior allele frequency (M*_Pstd_*, BayPass core-model output) at each outlier locus, identified the two populations showing the greatest deviation from zero, and computed their ratio *r* as a measure of signal concentration; outliers with *r* above the 80th percentile of the ratio distribution were classed as population-specific. For genotype–environment associations (GEA), we used an RDA (vegan v2.7-5; Oksanen *et al*. 2026) in addition to the BayPass auxiliary covariate model (MCMC sampling; Gautier 2015), since RDA explicitly partitions shared variance among correlated predictors, whereas BayPass treats covariates as independent within a joint SNP-level model, so correlated predictors can still inflate false-positive rates for individual covariates under this approach (Forester *et al*. 2018; Rellstab *et al*. 2015). We defined final GEA candidates as the union rather than the intersection of SNPs detected by either method, to avoid the higher false-negative rate of intersection-based approaches (Forester *et al*. 2018), while treating convergent detection as an indicator of higher confidence. Of nine initial environmental predictors, collinear variables were removed based on ecological relevance (latitude vs. distance from the Canche; elevation vs. river length), leaving five predictors with negligible residual multicollinearity (variance inflation factors). BayPass auxiliary-covariate runs used the same MCMC settings and significance calibration as the XtX analysis; candidates exceeded either the 99.5th POD percentile or the classical Jeffreys’ threshold (BF*_dB_* > 20). We performed a standard RDA (adjusted R², permutation-based ANOVA, 999 permutations) and a partial RDA (pRDA) conditioning on the first three principal components of Ω to control for neutral demographic covariance, identifying candidates as loci with loadings beyond ±3 standard deviations on significant constrained axes (Forester *et al*. 2018) and assigning each candidate to its most correlated predictor based on Pearson correlations. Finally, we cross-referenced predictor-specific GEA candidates against the five C2 regional outlier sets, reporting the region showing the largest numerical overlap for each predictor, to test whether environmental associations were diffuse or concentrated in particular metapopulations.

## Results

### Population structure, genetic diversity and effective population size

PCA revealed clear spatial structuring across the first three axes: PC1 and PC2 together explained 10.1% of genomic variance (5.7% and 4.4%, respectively; Fig. S4), with PC3 adding 1.5% (Fig. S5). PC1 separated Upper Normandy, Touques and Frome from all other populations, while PC2 further distinguished Brittany and Lower Normandy from the remaining groups. In PC1–PC3 space, Allier (ALL) occupied an extreme position and Southwest Atlantic rivers formed a distinct cluster at the opposite end of the axis. Clustering analyses supported this hierarchical structure: DAPC (BIC) and ADMIXTURE (cross-validation error) both identified five main genetic clusters (Fig. 1, S6, S8): Southwest (GAV, NIV, NVL, BID), Allier (ALL), Brittany (ELL, SCO, BLA, AUL, TRI), Lower Normandy/Mont-Saint-Michel Bay (SIE, SEE, SEL, COU, TOU), and Upper Normandy/English Channel (ARQ, BRE, CAN, FRO). Southwest and Upper Normandy populations were the least admixed; Trieux and Couesnon showed mixed Brittany/Lower Normandy ancestry, while Tamar and Orne did not align with any single group. RDA constrained by river-scale environmental predictors recovered the same five clusters (Fig. 2).

**Figure 1.**
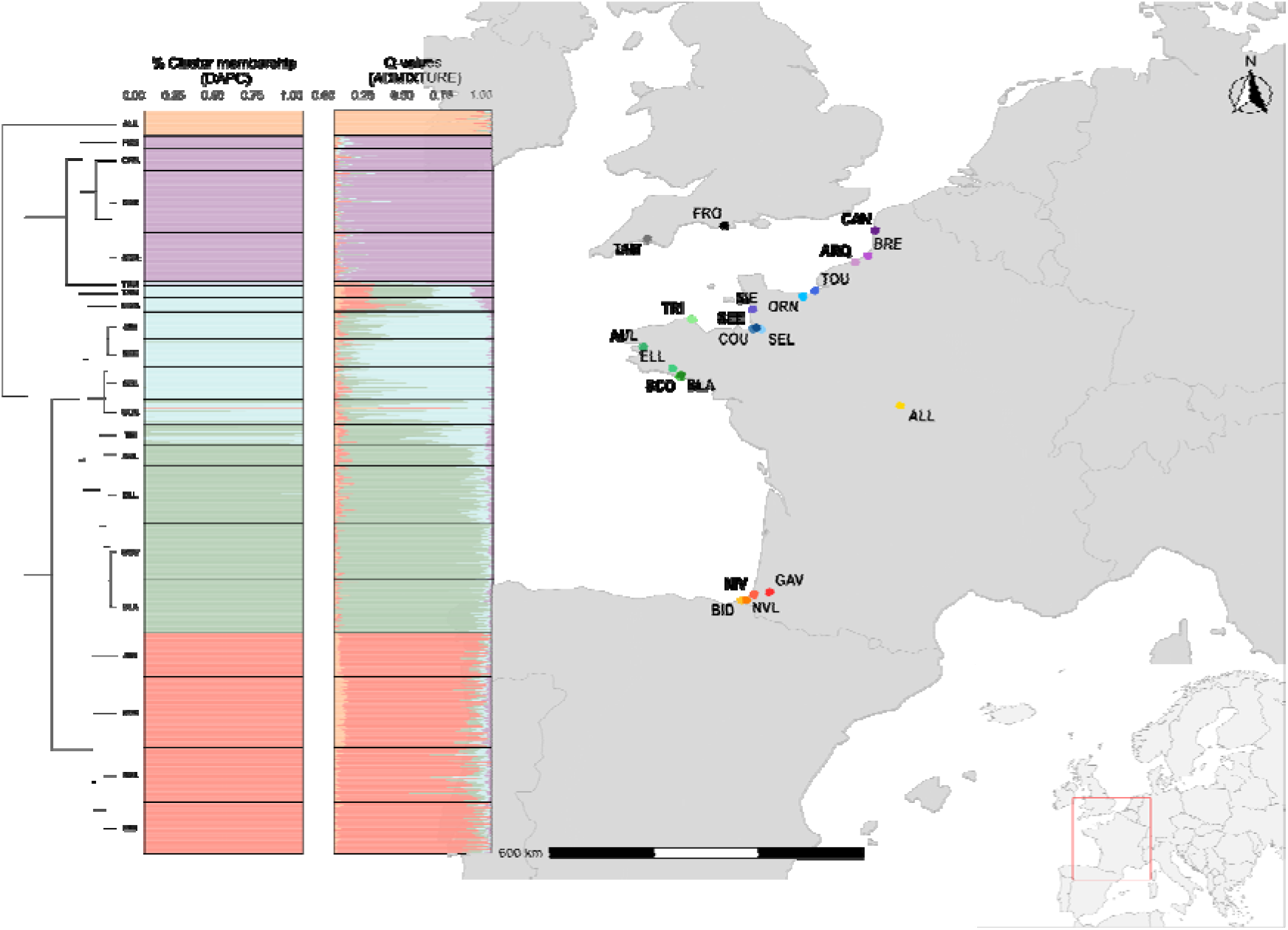
Neutral population structure of Atlantic salmon across sampled rivers. Left: Nei’s dendrogram based on pairwise genetic distances among the 21 sampled populations; only nodes supported by ≥95% of bootstrap replicates are labelled. Centre: cluster membership proportions from Discriminant Analysis of Principal Components (DAPC, left column) and admixture proportions (Q-values) from ADMIXTURE (right column), both computed for K = 5 genetic clusters; each row represents one individual, grouped and labelled by population. Right: geographic map of the 21 sampling sites along the Bay of Biscay and the English Channel, with each point coloured by population; scale bar (600 km) and north arrow indicate spatial scale and orientation. Inset (bottom right): location of the study area (red box) within Europe.

**Figure 2.**
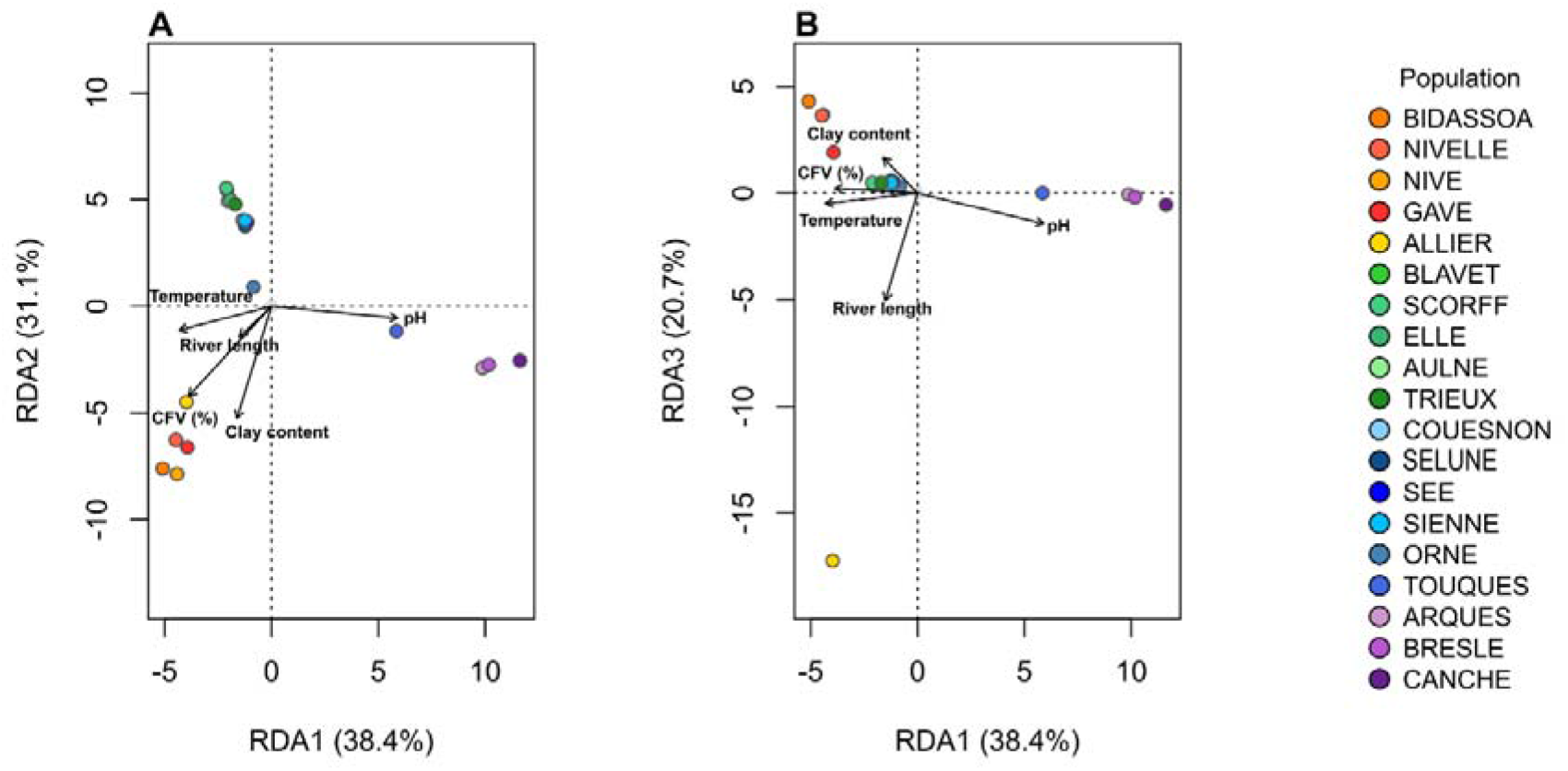
Redundancy analysis (RDA) of genomic variation constrained by environmental variables. Biplots of the first canonical RDA axis against RDA2 (panel A) and RDA3 (panel B), obtained from redundancy analysis of population-level allele frequencies constrained by river-scale environmental descriptors. Points represent river populations coloured by genetic clusters, ordered from south to north. Arrows indicate standardized canonical loadings of environmental covariates (mean water temperature, pH, clay content, coarse fragment volumetric (CFV%), and river length).

Pairwise *F*_ST_ followed a clear hierarchical pattern, with lowest values within clusters and highest among distant clusters (Fig. S1), and Allier showing the highest values overall (up to *F*_ST_ = 0.196 for the ALL–CAN comparison). Hierarchical AMOVA (Allier excluded) confirmed significant differentiation at both levels, with variance among groups (Φ = 0.137, p < 0.001), roughly five times that among populations within groups (Φ = 0.029, p < 0.001), yielding an overall Φ of 0.162 (p < 0.001). Mean expected heterozygosity ranged from 0.31 (Sée) to 0.39 (Nive), matching closely observed heterozygosity across most populations. Private alleles were concentrated in the Southwest populations. *F_IS_* values were low and sometimes significant but always < 0.01 (Table 1).

Contemporary Ne varied widely across the three CurrentNe2 models (Fig. S2, Table S2): under inter-chromosomal panmixia, Ne ranged from 75.2 (Orne) to 1726.4 (Ellé), with Gave, Tamar and Sienne also showing high Ne values. In five rivers — Scorff in Brittany, Orne and Sélune in Lower Normandy, and Nivelle and Bidasoa in the Southwest — Ne estimates under the subdivision-with-migration model significantly exceeded the inter-chromosomal panmictic estimate (non-overlapping 90% CIs; Table S2), consistent with these rivers functioning as connected patches within a larger metapopulation. Conversely, in eight rivers, including the least admixed and most differentiated populations in the dataset (Ellé, Gave, Nive, Allier), the subdivision estimate was significantly lower than the panmictic one — a direction opposite to that expected under population subdivision (Santiago et al., 2025).

### Identification of dispersers and population assignment tests

Cross-validation accuracy differed markedly among geographic groups (Table S3; Fig. S11– S20): highest in the Southwest and Upper Normandy (Monte-Carlo accuracies 0.88–0.90 and 0.89–0.92), intermediate in Brittany (0.79–0.88), and lowest in Lower Normandy (0.55–0.69), consistent with weaker within-cluster differentiation there. Five-fold results were consistent (Southwest 0.89, Brittany 0.81, Lower Normandy 0.64, Upper Normandy 0.91; global baseline 0.84–0.86).

Individual assignments showed a clear contrast between among- and within-cluster resolution (Fig. 3): adults were assigned homogeneously within clusters, with abrupt transitions between them, except Trieux and Couesnon which showed mixed Brittany/Lower Normandy assignment. Within clusters, the proportion of unassigned adults (q < 0.70) increased from the Southwest through Brittany and Upper Normandy to a maximum in Lower Normandy (91.3–100%). In the Southwest (n = 225 adults), assignment remained well resolved: Gave and Nive had the highest local assignment (89.2% and 89.1%; 10.8% and 6.5% unassigned). Putative dispersers were detected in Nivelle (11.5%), Bidasoa (9.4%) and Nive (4.4%) but not Gave. In Brittany (n = 169), resolution was lower (43.9–72.9% unassigned); Ellé and Scorff retained more local adults (56.1%, 32.1%) whereas Blavet had only 16.9% of local adults, though the high unassigned proportion precludes a reliable dispersal estimate. In Lower Normandy, unassigned proportions were extremely high in all four rivers (91.3–100%), precluding any dispersal estimate. Upper Normandy combined the highest assignment accuracy with low local-assignment proportions (18.2% Arques, 31.7% Bresle); Canche showed an extreme sink pattern (0% locally assigned, 63.2% unassigned), so these figures represent a lower bound.

**Figure 3.**
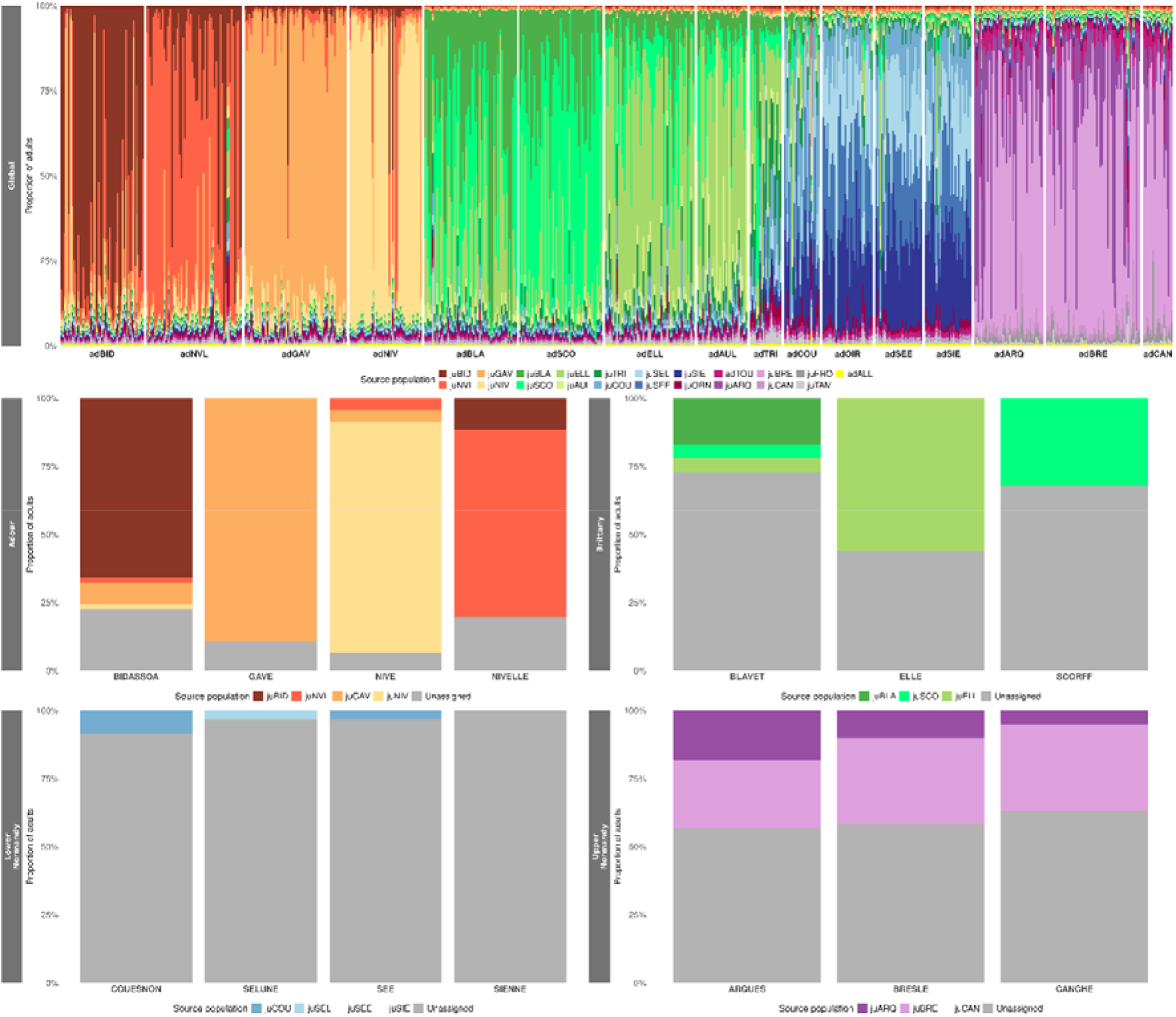
Population assignment of adult Atlantic salmon. Top: individual membership probabilities from the global assignment framework, showing each adult’s full probability distribution across juvenile source populations. Bottom: River-level dispersal rates within four regional metapopulations (Southwest, Brittany, Lower Normandy, Upper Normandy). Bars show the proportion of adults confidently assigned to each source river (q ≥ 0.70); individuals below this threshold were classified as unassigned (grey).

### Adaptive differentiation

The global XtX genome scan identified 132 outlier SNPs (Fig. 4B; Table S5A), unevenly distributed across 24 chromosomes: Ssa13 alone held 28 outliers, followed by Ssa1 (15 outliers), Ssa9 (14 outliers) and Ssa14 (13 outliers), while most chromosomes had 1 to 8. Outlier XtX values ranged from 31.9 to 49.5. The strongest signals included a Ssa19 SNP overlapping *vps41* (lysosomal trafficking/autophagy) and a Ssa13 SNP overlapping the MUC18-like gene *LOC106567672* (cell adhesion). Including putative dispersers reduced the outlier set to 41 SNPs (Fig. 4A; Table S5B); 92 outliers were detected only when dispersers were excluded, and just one was unique to the disperser-inclusive scan. The local-score approach identified four significant genomic windows (Fig. 4C): Ssa3 (82.9–83.6 Mb, 715.8 kb), Ssa9 (30.1 Mb, 44.8 kb) and two on Ssa13 (31.2–32.3 Mb, 1,051.2 kb; 56.7–57.1 Mb, 400.1 kb), containing 6–17 outlier SNPs and 4–13 genes each (10–67 genes/Mb; Table S6A). The Ssa9 window was exceptionally gene-dense (∼67 genes/Mb, over three times the genome average; Lien et al. 2016). Peak SNPs overlapped *LOC123741732* (Ssa3; ER-to-Golgi vesicle transport), *ctage5* (Ssa9; ER-to-Golgi trafficking, trans-Golgi network), *LOC106566887* (Ssa13, 31–32 Mb; transcription factor) and *LOC106567491* (Ssa13, 57 Mb; phenylalanyl-tRNA ligase) (Table S6B). Ranking populations by M*_Pstd_* at each of the 132 global outliers showed uneven contributions (Fig. 4D): Nive (27), Allier (23) and Arques (18) were most often top-ranked, together accounting for 51% of attributions, whereas Blavet, Bresle and Aulne were rarely primary sources (Table S7A). Twenty-seven outliers (20%) were classified as population-specific (*r* > 1.78, 80th percentile), predominantly attributed to Allier (9) and Arques (7; 59% combined), followed by Nive (4); the remaining outliers showed more diffuse, shared differentiation.

**Figure 4.**
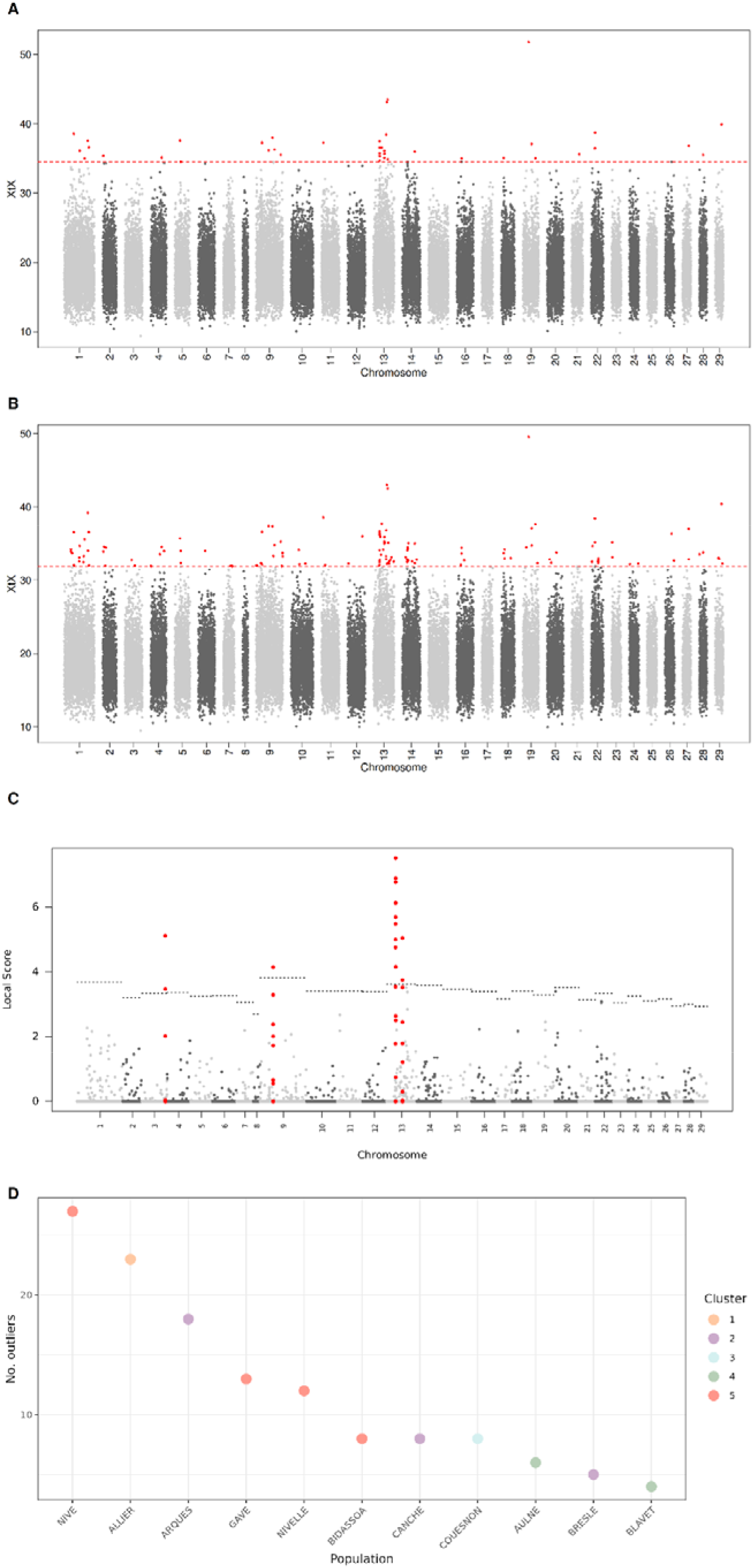
Genome-wide scans for adaptive differentiation, and population-specific contributions to the outlier signal. (A, B) Manhattan plots of XtX values across the 29 Atlantic salmon chromosomes, computed under the BayPass core model on the local-only dataset including putative dispersers (A) and excluding them (B); red points indicate outlier SNPs exceeding the 99.5% quantile of the calibrated empirical XtX null distribution (dashed red line). (C) Local-score statistics computed from XtX p-values (ξ = 2; Fariello et al. 2017) on the local-only dataset; red points represent contiguous genomic windows enriched in candidate SNPs, and dotted lines indicate the chromosome-specific significance threshold. (D) Number of outlier SNPs for which each population ranked as the top contributor to the differentiation signal: for each of the 132 outlier SNPs, the population with the highest absolute standardized posterior mean allele frequency (M_Pstd) was identified as the primary source of the signal.

Within-cluster XtX genome scans consistently identified outliers despite weaker neutral differentiation among populations than among clusters, with candidate sets ranging from 145 SNPs in Lower Normandy to 214 in the Southwest (Fig. 5; Table S8). POD-calibrated thresholds (99.5th percentile) varied moderately across clusters (Southwest: 12.03; Lower Normandy: 11.97; Brittany: 10.31; Upper Normandy: 8.81), and maximum XtX values within clusters ranged from 17.5 to 21, compared to 31.9–49.5 at the global scale. Overlap with the 132 global outliers (Fig. S21) ranged from 3 SNPs (Lower Normandy, 2%) to 49 (Southwest, 37%), with Upper Normandy and Brittany intermediate (12 each, 9%) — an asymmetry not explained by differences in outlier-set size (Lower Normandy n = 145 vs. Southwest n = 214).

**Figure 5.**
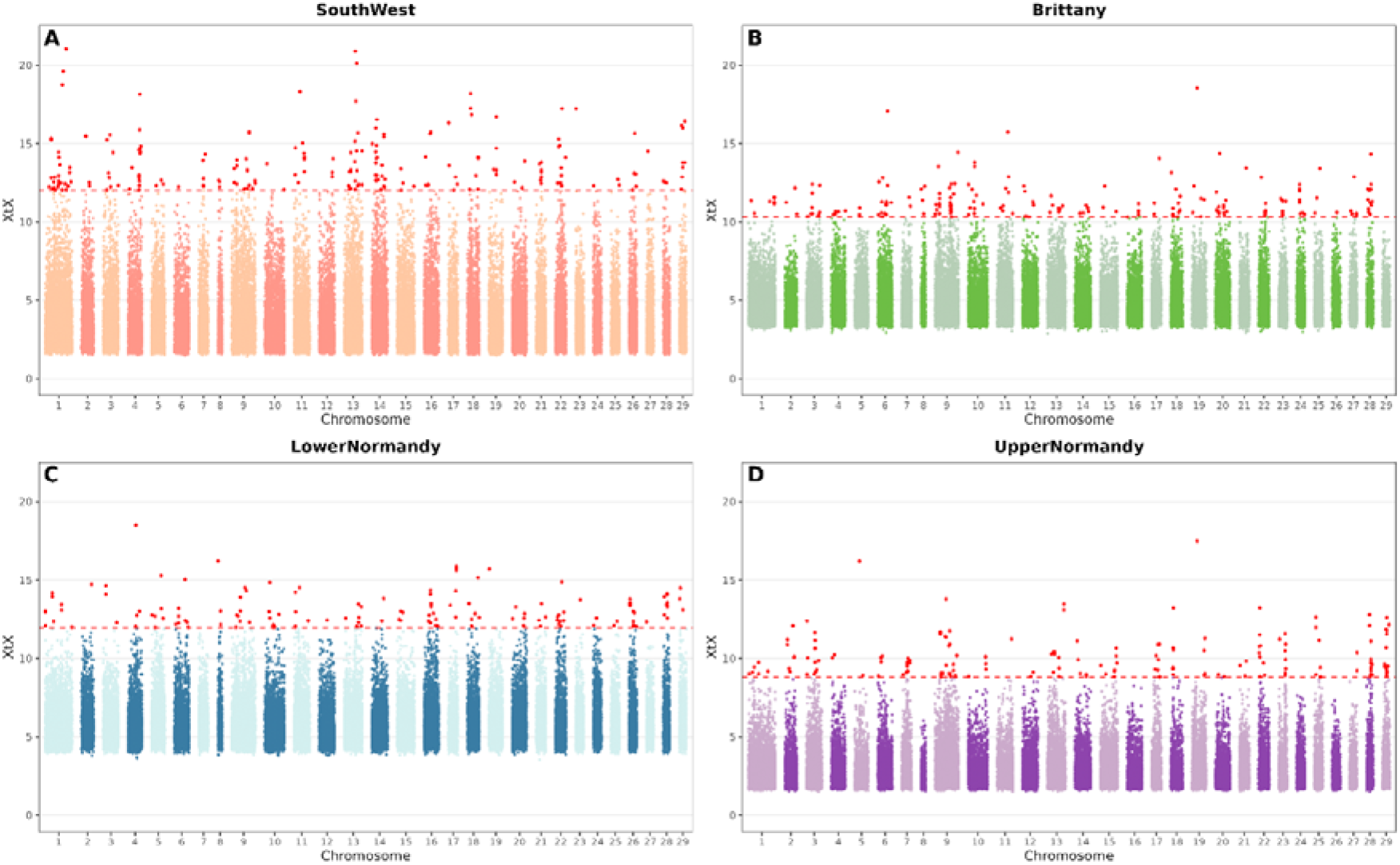
Within-cluster genome scans. Manhattan plots show XtX values for SNPs across the genome within each regional cluster after exclusion of putative dispersers: (A) SouthWest, (B) Brittany, (C) Lower Normandy, and (D) Upper Normandy. Red points indicate SNPs exceeding the metapopulation-specific 99.5% POD-calibrated XtX threshold, shown by the red dashed horizontal line.

C2 contrasts identified 233 (Brittany, threshold 6.54) to 575 (Allier, threshold 3.49) outliers, with Lower Normandy (315), Upper Normandy (283) and Southwest (251) intermediate (Fig. 6; Table S9). Pairwise overlap among C2 sets was limited (Fig. S22), largest for Allier (21 shared with Southwest, 20 with Lower Normandy). Overlap between these C2 outliers and the global XtX outliers ranged from 4 (Brittany, 3%) to 30 SNPs (Allier, 23%), with Southwest, Lower Normandy and Upper Normandy intermediate (7, 9, 7; 5–7%).

**Figure 6.**
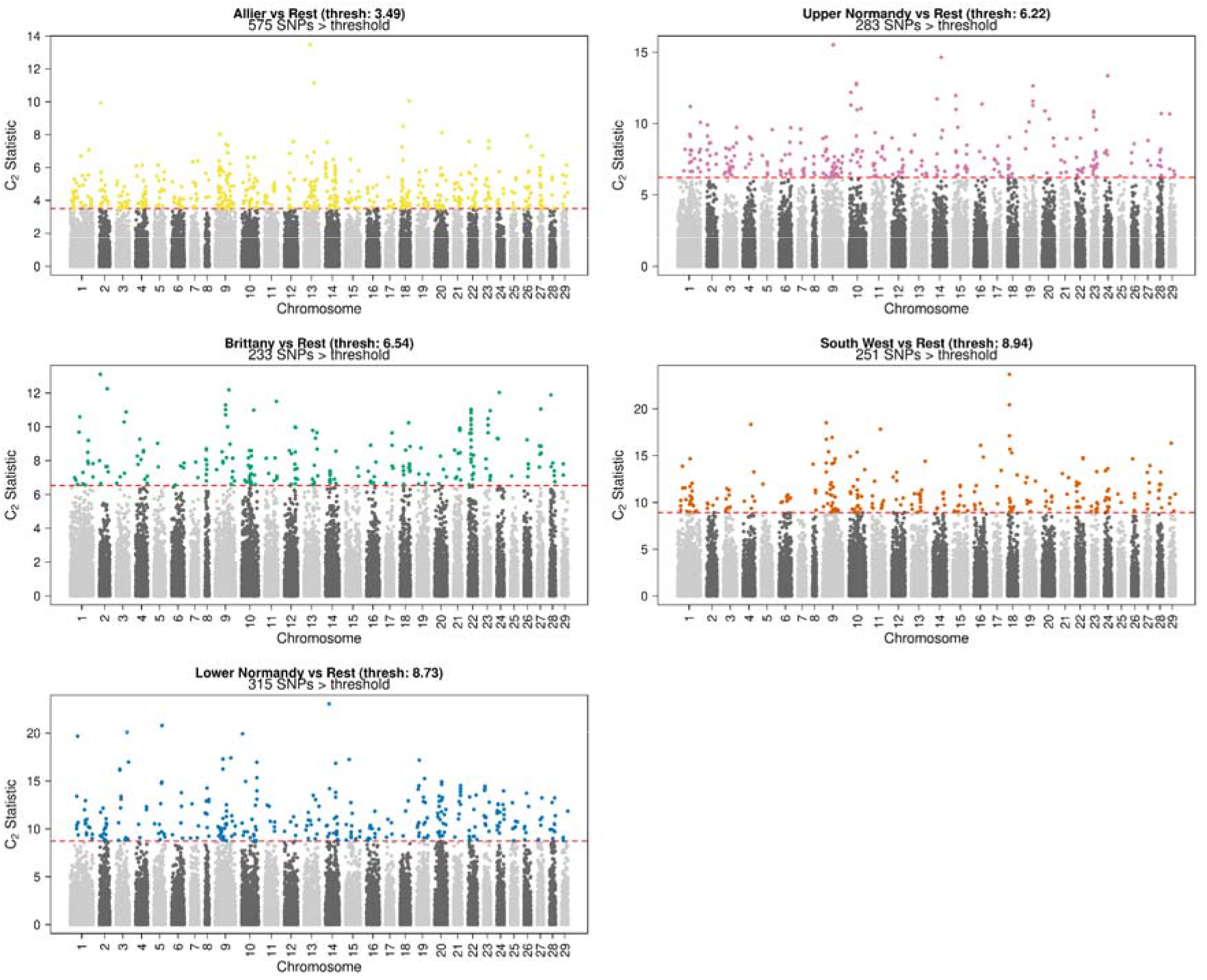
Genome-wide C2 contrast statistics for five one-vs-rest regional comparisons. Manhattan plots showing the C2 statistic (Y-axis) across the 29 Atlantic salmon chromosomes (X-axis) for each of five contrasts: Allier vs. Rest of France, Upper Normandy vs. Rest, Brittany vs. Rest, Southwest vs. Rest, and Lower Normandy vs. Rest, computed under the BayPass core model on the local-only dataset. Red dashed lines indicate the contrast-specific 99.5th percentile threshold derived from the pseudo-observed dataset (POD) null distribution. SNPs exceeding the threshold (outliers) are colour-coded by the region. The number of outlier SNPs per contrast is indicated above each panel.

### Genome–environment associations

The five retained environmental predictors jointly explained substantial genomic variance in the RDA (R²_adj_ = 0.46; R² = 0.61; permutation test: F, = 4.11, p = 0.001), with three significant constrained axes accounting for 90.1% of constrained variance (RDA1: F = 7.88; RDA2: F = 6.88; RDA3: F = 4.91; all p = 0.001; RDA4–5 non-significant). RDA1 separated Upper Normandy/Picardy populations (Touques, Arques, Bresle, Canche) along a pH gradient; RDA2 distinguished cooler Brittany/Lower Normandy rivers from warmer Southwest rivers (Fig. 2); RDA3 (20.7% of constrained variance) isolated Allier along the river-length gradient — an axis structure that closely mirrors the neutral genetic clusters. A ± 3 SD loading threshold on the pRDA axes identified 552 candidates (Table S10): pH (159), river length (109), clay content (103), temperature (91) and CFV(%) (90). BayPass POD-based candidates ranged from 96 (river length) to 346 (temperature) (Fig. 7; Table S11), and BF > 20 candidates from 13 (clay content) to 32 (pH); distributions were largely diffuse except for concentrated clusters on Ssa13 (river length, clay content), Ssa14 (CFV%) andSsa9 (pH).

**Figure 7.**
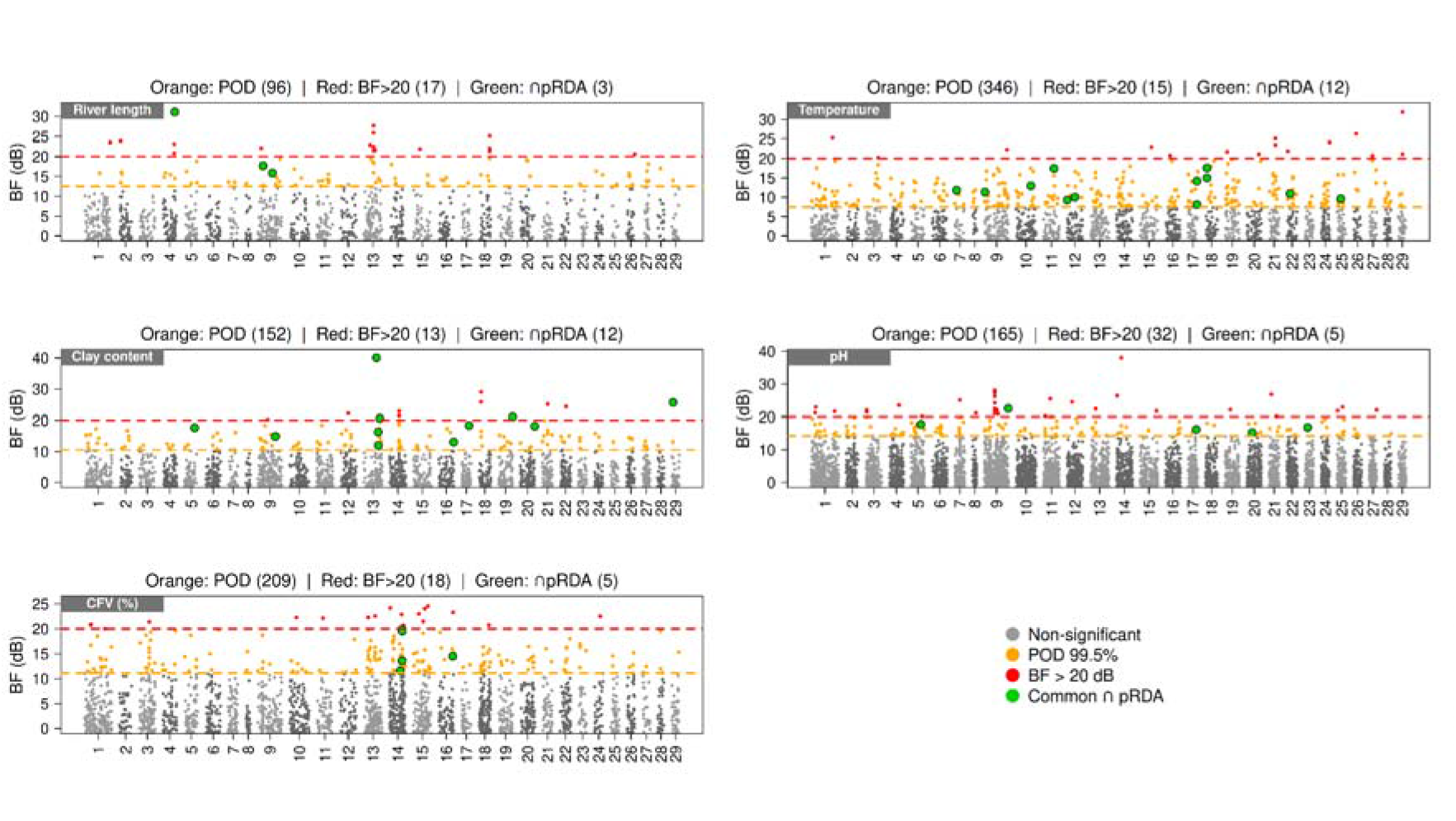
Genome-wide Bayes Factor scans for genotype–environment associations with five environmental predictors. Manhattan plots of Bayes Factors (BF, in deciban units) obtained from the BayPass Standard covariate model for river length, temperature, clay content, pH, and coarse fragment volumetric content (CFV, %). Orange points indicate SNPs exceeding the 99.5% quantile of the POD-simulated null BF distribution; red points indicate SNPs exceeding the BF > 20 dB threshold (Gautier 2015). Green-outlined points indicate candidate SNPs also identified by the partial RDA (pRDA) for the corresponding predictor. The number of candidates detected under each criterion (POD, BF > 20, and pRDA) is indicated above each panel.

Thirty-seven SNPs were detected convergently by pRDA and BayPass (Fig. 7; Table S12), most strongly for temperature and clay content (12 each), moderately for pH and CFV% (5 each), and least for river length (3). Four of these 37 GEA candidates were also XtX outliers (triple-overlap set; Table S12B): clay content (Ssa13, near *ints2*; XtX = 42.5), pH (Ssa9, intronic in *LOC106613081*; XtX = 33.7), and river length (Ssa4, near *tmem132e*, XtX = 33.9; Ssa9, near *ctage5*, XtX = 32.1 — overlapping the Ssa9 local-score window with its highest XtX value). Cross-referencing predictor-specific GEA outliers against the five C2 sets showed uneven regional concentration. River length stood out: 52% of BayPass and 15% of RDA river-length candidates were also Allier C2 outliers, by far the largest overlap of any predictor–region pair. pH candidates overlapped most with Upper Normandy under BayPass (10%), consistent with the pH-driven RDA1 axis, though weak and diffuse under RDA. Clay content and temperature showed no consistent regional concentration between methods, and CFV (%) overlap was weak throughout (≤5 SNPs in any contrast).

## Discussion

At the southern edge of its European range, we show that the hierarchical structure of Atlantic salmon populations is associated with genome-wide signatures of putative adaptive differentiation at two spatial scales: among clusters and among neighbouring rivers connected by contemporary dispersal. These signatures are not distributed uniformly across populations: they remain sharp in some rivers despite immigration, and are already blurred in others where gene flow has homogenised the genetic background. Together, these findings demonstrate the interplay of dispersal and adaptation at different spatial scales in southern Atlantic salmon populations of high conservation concern (Gabián *et al*. 2022; Valiente *et al*. 2010).

### The hierarchical genetic structure provides the demographic context for genome-wide inference

The delineation of five major genetic groups — Southwest, Brittany, Lower Normandy/Mont-Saint-Michel Bay, Upper Normandy/English Channel and Allier — with differentiation among clusters (AMOVA Φ = 0.137) roughly five times higher than among rivers within clusters (Φ = 0.029), confirms with genome-wide markers the hierarchical structure described from microsatellites by Perrier et al. (2011). Contemporary effective-size estimates support this reading. In five rivers spanning three clusters — Scorff in Brittany, Orne and Sélune in Lower Normandy, and Nivelle and Bidasoa in the Southwest — Ne under the subdivision-with-migration model significantly exceeded the inter-chromosomal panmictic estimate, indicating that these rivers are best treated as connected patches within a larger metapopulation rather than as demographically closed units (Ryman & Laikre 1991; Santiago *et al*. 2025). Rivers within a metapopulation share recent ancestry and exchange spawners within a single generation, whereas metapopulations are separated by broader climatic and geological contrasts and by much lower effective gene flow. Scans run at both levels therefore address different questions, and there is no reason to expect them to return the same outliers. Their results are also not interchangeable: because XtX and C2 are calibrated against the covariance matrix Ω of the populations included in each analysis (Gautier 2015), outlier magnitudes and thresholds are not comparable across scans, and we restrict between-scale comparisons to the identity and genomic position of candidate SNPs.

### From dispersal to gene flow

Rather than a single connectivity regime, we find four contrasting configurations, in which the level of differentiation among neighbouring rivers governs both how much dispersal occurs and how much of it assignment can resolve. In the Southwest, where differentiation among nearby rivers is highest, assignment resolution was accordingly the highest: putative dispersers were detected in three of the four rivers while most adults remained confidently local. For one river, an independent fitness-based estimate of origin-dependent dispersal is available (Egal *et al*. 2026), and our results converge with it for two of its three neighbouring sources. Nive, the most differentiated source in our genome-wide estimates, contributed no detectable dispersers in our independent sample, consistent with the severe reproductive deficit reported for Nive-origin immigrants in that system. Conversely, the least differentiated source accounted for essentially all dispersers we identified, mirroring its earlier documentation as the dominant source of dispersal. Only the third, intermediate source diverges between datasets, a discrepancy more plausibly attributable to differences in sampling depth and temporal coverage than to a true absence of connectivity. Nive is therefore an informative case: it sends almost no detectable immigrants into the neighbouring river, the few Nive-origin immigrants recorded there reproduced poorly, and it is the population most often carrying the most extreme standardised allele frequency across the national outlier set. Whether limited exchange allowed that divergence to accumulate, or whether divergence now penalises Nive-origin immigrants elsewhere, cannot be separated with these data; both act in the same direction and at successive stages, leaving the alleles that distinguish Nive little opportunity to enter any neighbouring gene pool.

In Brittany and Upper Normandy, where local gene flow was much higher, local dispersal rates could not be inferred; considering only identified dispersers and locals, however, the relative origins of dispersers remain informative. In Brittany, Ellé and Scorff retain most local adults and supply the dispersers found in Blavet, a net recipient; in Upper Normandy, Arques and Bresle exchange migrants reciprocally, while Canche receives immigrants from both, a putative sink.

We use source and sink in the net-flow sense of Loreau *et al*. 2013— net exporter or importer of assigned adults within our sampling windows — not in the demographic sense. Moreover, a net flow of individuals is not a measure of genetic contribution, since a net recipient may still be a substantial genetic contributor. Such asymmetric connectivity in a metapopulation can buffer demographic fluctuations and promote genetic rescue (Lamarins *et al*. 2022, 2024b), or erode local adaptation if gene flow outpaces selection (Kawecki & Holt 2002; Lenormand 2002); the outcome will depend on whether selection in these rivers is stable or fluctuating over time (Blanquart *et al*. 2013). Lower Normandy could not be placed on the same footing, since the low resolution of its baseline prevented the identification of dispersers altogether. The documented history of stocking in that area could contribute to this pattern (Perrier *et al*. 2013). Interestingly, Trieux, the geographically closest population between Brittany and Upper Normandy, seems to reveal a contact zone between those two clusters given the level of local admixture, which could also be explained by stocking history. The global pattern remains consistent with strong philopatry and with a dispersal propensity declining with distance (Birnie-Gauvin *et al*. 2019; Hasler & Scholz 1983; Lamarins *et al*. 2024a; Salmenkova 2017; Stabell 1984). Which individuals disperse, and where, may depend on sea age, the spatial configuration and environmental similarity of neighbouring rivers, local abundance and the demographic balance between source and recipient populations (Chat *et al*. 2022; Hamann & Kennedy 2012; Jonsson *et al*. 2003; Westley *et al*. 2025). Except in the Southwest, where origin-specific fitness data point to selection against immigrants (Egal *et al*. 2026), our data cannot identify which mechanism underlies the pattern.

Whether dispersal translates into gene flow is a separate question, because gene flow additionally requires that dispersers survive, reproduce, and produce viable offspring. Dispersal is therefore only an upper bound on realised gene flow, and the gap can be wide: in two species sharing a landscape, indistinguishable dispersal rates translated into a six-fold difference in gene flow, driven by effective size rather than by behaviour (Bouchard *et al*. 2025). Part of that gap can also be methodological, since some adults scored as dispersers may represent temporary straying rather than settled recruitment (Keefer & Caudill 2014). In our own data, including putative dispersers in the XtX scan reduced the candidate set from 132 to 41 SNPs; 92 candidates were detected only once dispersers were removed, meaning non-local genotypes measurably attenuate allele-frequency contrasts among rivers even within a single generation of sampling. Whether this attenuation reflects an ongoing homogenising process or simple statistical dilution from pooling local and non-local genotypes depends entirely on whether those immigrants reproduce and recruit, which our design cannot test directly. Arques is the clearest case in the dataset where dispersal and gene flow visibly come apart. It has the most disperser-dominated assignment profile of any river, with fewer adults confidently local than confidently immigrant from Bresle, yet it is among the three populations most often carrying the extreme standardised allele frequencies at global outlier loci, and it hosts more population-specific outliers than any river except Allier. Consistent with this, the LD-based subdivision model gives no support for subdivision in Arques. If these immigrants were reproducing and recruiting at rates comparable to residents, contrasts at these loci should already be eroding.

The persistence of such contrasts is compatible with several non-exclusive explanations, two of which are known to sustain divergence within a dispersal neighbourhood (Richardson *et al*. 2014): post-arrival selection filtering non-local genotypes before their alleles enter the local gene pool (Nosil *et al*. 2005), and non-random dispersal, whereby immigrants are not a representative allele-frequency sample of their source at these loci (Edelaar & Bolnick 2012); a third is simply that immigration is too recent to have left a genomic signature. A fourth is architectural. Because assignment is based on the full, predominantly neutral SNP panel, weak genome-wide resolution and sharp divergence at a handful of loci are not in tension: this is what is expected if a subset of candidate regions behaves as semipermeable to gene flow while the rest of the genome introgresses more freely (Nosil *et al*. 2009), as demonstrated in fishes, where chromosomal rearrangements and low-recombination regions shield adaptive variants from homogenising gene flow (Barth *et al*. 2017; Lehnert *et al*. 2020; Tigano & Friesen 2016). Under this reading, Arques’ high proportion of unassigned adults would be a signature of genome-wide blending rather than statistical noise, and the loci retaining a clean local signal would be predicted to co-locate with regions of restricted recombination — plausibly the windows on chromosomes 3, 9 and 13.

### Adaptive differentiation occurs at both global and local scales

The genomic distribution of the 132 global genome scan outliers was markedly uneven: four local-score windows, on chromosomes 3, 9 and 13, concentrated a large share of the signal, with Ssa13 alone carrying 28 outliers (21% of the total). This concentration could simply reflect statistical non-independence: SNPs in physical linkage with one or a few causal variants co-segregate and are jointly flagged, so the number of independent targets is smaller than 132 outliers suggest. Local variation in recombination amplifies this, since differentiation is higher and false positives more frequent where recombination is low: in coho salmon, both differentiation scans and GEAs were significantly affected by recombination-rate variation (Rougemont *et al*. 2023). We cannot correct for this confounder, which requires a recombination map that an array-based design does not provide. Recombination is strongly localised along the Atlantic salmon genome (Lien *et al*. 2011), where structural variants can generate discrete linkage blocks behaving as single units (Bertolotti *et al*. 2020). A second possibility is that the target is genuinely gene-dense, as illustrated by the Ssa09 local-score window, which at ∼67 genes/Mb is more than three times the genome-wide average (Lien *et al*. 2016) and could plausibly harbour several linked, independently favoured genes rather than one. A similar concentration of selection and GEA signals on Ssa09, on a gene-rich haploblock containing six6, has been reported at the within-population scale in a Baltic system (Miettinen *et al*. 2023). A third possibility is that these windows are just coincidences of a broader differentiation landscape. Large trans-Atlantic differentiated regions have been reported on Ssa13 and Ssa19 — two of the chromosomes carrying our strongest signals — and, importantly, showed elevated linkage disequilibrium without any detectable reduction in recombination (Lehnert *et al*. 2020), which cautions against treating low recombination as the default explanation. We cannot arbitrate among these without finer-scale recombination and haplotype data. What can be said is that all three readings remain compatible with theory predicting that adaptive architecture under migration–selection balance evolves toward concentration in few, tightly linked regions, precisely because such architectures resist swamping more effectively than a diffuse polygenic basis (Yeaman & Whitlock 2011).

In addition, independent within-cluster scans detected outlier sets of the same order across the four regions, confirming that adaptive differentiation is not restricted to broad regional contrasts but is also present among neighbouring rivers experiencing contemporary dispersal (Brauer *et al*. 2016). Their overlap with the global list is partial, and its unevenness is informative at the extremes. The Southwest, where differentiation among populations is highest, shows the strongest coupling between global and local outlier sets, sharing 37% of the global outliers: the same loci differentiate its populations from each other and from the rest of France, and Nive, itself receiving a small share of dispersers, is nonetheless the population most often carrying the extreme standardised allele frequency at global outlier loci. Lower Normandy, where differentiation among rivers is lowest, shares 2% and is the weakest contributor to every outlier statistic computed here, though this must be read cautiously since 91–100% of its adults fell below the assignment confidence threshold and were nonetheless retained in the genome scans. Both extremes converge on the same reading: outliers are detectable where allele-frequency contrasts among populations remain large, and undetectable where gene flow — natural, or linked to stocking history (Perrier *et al*. 2013) — has already homogenised the genetic background (Lenormand 2002). The four scans were run on comparable numbers of populations, so the contrast is unlikely to reflect differences in the number of sampled demes, a factor known to inflate outlier detection in river networks (Fourcade *et al*. 2013). Arques makes the same point at the scale of a single river: what matters is not the dispersal rate but whether it has translated into genome-wide gene flow.

At the genetic cluster scale, pairwise overlap among the five C2 outlier sets was limited beyond the comparatively large intersections involving Allier, indicating that each cluster’s distinctiveness is carried largely by a different subset of loci rather than a shared, recurrently used genomic toolkit. This is consistent with a broader expectation that replicate populations facing different combinations of environmental contrast and connectivity rarely converge on identical genomic solutions, even to broadly similar selective problems (Bolnick *et al*. 2018). Whether this reflects genuinely different selective pressures or a shared pressure met by different genetic solutions within a polygenic, redundant architecture (Bernatchez 2016; Pritchard & Di Rienzo 2010) cannot be resolved from outlier identity alone: it would require comparing functional annotation rather than identity across regions, a natural extension of this dataset.

One caveat is specific to the within-cluster scale. Because neutral differentiation among populations within clusters is much lower than among clusters, within-cluster models are calibrated on a narrower range of covariance, which can allow more SNPs to exceed a given quantile without any genuine increase in selection strength (Gautier 2015). We therefore treat within-metapopulation outlier sets as evidence that fine-scale differentiation exists, but not as a quantitative measure of relative selection strength. More generally, outlier loci may reflect divergent selection, but also linked selection, recombination-rate heterogeneity, structural variation or residual demographic effects (Bierne *et al*. 2011; Hoban *et al*. 2016; Lehnert *et al*. 2020; Roesti *et al*. 2012).

### Environmental associations and candidate loci

The environmental association results point to a multivariate rather than a single selective axis. Climatic, geological and hydrographic variables jointly explained a substantial share of genomic variance, consistent with other salmonid and riverine landscape-genomic studies in which multivariate gradients rather than single predictors best explain adaptive divergence (Bourret *et al*. 2013; Brauer *et al*. 2016; Dallaire *et al*. 2026; Hecht *et al*. 2015). As in those systems, the three significant constrained axes closely mirror the neutral genetic clusters, underscoring a covariance among environment, geography and demographic history that population-level data alone cannot fully partition (Bekkevold *et al*. 2020; Moore *et al*. 2014). This covariance is amplified with the cross-referencing of GEA outliers against the regional C2 sets, where for four of the five predictors — pH, clay content, temperature and CFV(%) — regional concentration was weak, as expected under diffuse multivariate selection. River length appears as an exception: more than half of the BayPass GEA candidates were attributed to C2 outliers, with a smaller but concordant fraction under RDA. That concentration is coherent with everything else we observe for Allier. Its C2 contrast yielded more candidates than any other regional comparison, its overlap with the global XtX list is the largest of the five, RDA3 isolates it along the river-length gradient alone, and it shows no admixture in the clustering results (Perrier *et al*. 2011, 2013). Allier adults complete the longest anadromous migration recorded in western Europe, 750–950 km to the spawning grounds, almost exclusively as multi-sea-winter fish (Evanno *et al*. 2023). This river-length signal seems to be attributable to selection associated with migration distance itself — energetic allocation, return timing, sea-age structure — concentrated in one exceptional population. The same signal was recovered by Rougemont *et al*. (2023) at continental scale in coho salmon, and after explicitly controlling for the recombination confounder discussed above, migration distance emerged as the primary selective factor driving local adaptation and partial parallel divergence among distant populations. It is further corroborated by common-garden evidence that Allier embryos differ from other French populations in their reaction norms to hypoxic stress (Côte *et al*. 2012) and in embryonic thermal plasticity (Côte *et al*. 2016). Our genome scan adds a genomic line of evidence to a pattern already established at the phenotypic level, and provides a first set of positional candidates.

The second most concentrated predictor, pH, points to geology-driven water chemistry, and is the axis along which RDA1 separates the Upper Normandy and one Picardy river from the rest. King *et al*. 2026 independently recovered the same chalk-versus-non-chalk contrast — driven by temperature and precipitation variables tied to the underlying geology — as the primary axis separating Upper Normandy and the chalk rivers of southern England from neighbouring non-chalk populations. The regional cross-referencing supports this only partially, pH candidates overlapping most with the Upper Normandy C2 set under BayPass but weakly and diffusely under RDA.

Only 37 SNPs were detected convergently by both GEA methods, four of which were also national XtX outliers. In this triple-overlap set, where our lines of evidence coincide most closely, one locus stands out. The Ssa09 SNP near ctage5 falls within the densest of our local-score windows, is the SNP with the highest XtX value in that window, and is associated with river length by both GEA methods. ctage5 participates in COPII-mediated ER-to-Golgi trafficking and, in mice, has been implicated in adipocyte differentiation, insulin signalling and lipolysis (Fan *et al*. 2025) — a coherent but untested link to energy balance, an axis closely tied to maturation timing in this species (Ahi *et al*. 2025). No direct evidence connects ctage5 to migration or maturation in Atlantic salmon, and we present it as a functional hypothesis rather than a mechanism. Ssa09 also carries a SNP near six6, previously linked to spawning-site selection and return-migration timing (Pritchard *et al*. 2018), and the pH-associated locus of the triple-overlap set; and it is the chromosome on which an independent Baltic study concentrated its own selection and GEA signals (Miettinen *et al*. 2023). Within Upper Normandy, we additionally recovered a locus near akap11, within the vgll3–akap11 maturation region on a different chromosome (Barson *et al*. 2015), together with one near negr1, associated with obesity and sexual maturity in other taxa (Lee *et al*. 2012). None of this is a demonstration since recurrent detection of a genomic region across studies does not establish that the same causal variant, haplotype or mechanism is involved in each system because at this marker density, and without haplotype-resolved data, our candidates locate regions rather than variants. Outliers near vps41, a MUC18-like locus, ints2 and tmem132e appear specific to this study, which may reflect genuine biological differences but also differences in marker density, reference genome or analytical design relative to previous work.

### Conservation implications: neither fully isolated nor fully interchangeable

Two main results could shape how these rivers are managed. First, exchanges among neighbouring rivers means local population dynamics is governed by connectivity within the broader network, not by local conditions alone (Lamarins *et al*. 2022), thus rivers cannot be treated as demographically closed units. Second, persistent fine-scale adaptive differentiation — even between rivers exchanging adults, as in Arques — argues against collapsing neighbouring rivers into a single management unit on connectivity grounds alone. Dispersal itself is neither uniformly beneficial nor costly: individual-based modelling shows a non-linear relationship between straying and metapopulation stability, intermediate rates maximising portfolio effects while very low and very high rates both erode robustness (Lamarins *et al*. 2022). Intermediate rates of gene flow could also maximise local adaptation under temporally fluctuating selection rather than just buffer demographic fluctuation, by replenishing genetic variation towards local optima (Blanquart *et al*. 2013). Maintaining connected river networks while preserving their environmental heterogeneity and regional distinctiveness is therefore a more defensible management objective than maximising either connectivity or isolation, particularly at the southern range edge, where declines have been steepest and climate-driven change in both freshwater and marine habitats is expected to be disproportionate (Almodóvar *et al*. 2019; Rikardsen *et al*. 2021).

The Southwest cluster deserves particular attention in this respect. It combines the strongest coupling between local and global candidate sets, the warmest thermal regime sampled, and a position at the species’ rear range edge. Rear-edge populations are increasingly recognised as reservoirs of adaptive potential, precisely because they already persist under conditions the rest of the range is only beginning to experience (Hampe & Petit 2005); the coherence of the Southwest signal across spatial scales strengthens the case for treating it as a conservation priority in its own right. Allier, conversely, illustrates the opposite conservation logic: an isolated, demographically fragile population (Evanno *et al*. 2023) whose adaptive distinctiveness — converging across phenotype, genome-wide differentiation and a single dominant environmental axis — is not a hypothesis but an established result, for which connectivity is not the relevant management lever. Where the Southwest argues for maintaining a connected network of environmentally distinct rivers, Allier argues for the opposite: protecting a single, irreplaceable, self-contained lineage. These two regions are therefore conservation priorities identified directly from the adaptive signal itself, rather than from neutral diversity alone. Importantly, our signal is drawn from the freshwater phase only and cannot capture selection or mortality during the marine phase that is now widely perceived as the principal driver of Atlantic salmon decline (Olmos *et al*. 2019, 2020).

## Conclusion

This study shows the interplay between dispersal and divergent selection in a hierarchical network, where dispersal does not necessarily translate into gene flow. Putative adaptive differentiation was strongest in a river where identified dispersers outnumbered identified locals, and weakest not where dispersal was highest, but where gene flow had already homogenised the genetic background beyond the resolution of assignment tests. Adaptive signals were detectable simultaneously at global, regional and river scales, and at each scale were concentrated in a small number of genomic regions rather than spread diffusely — consistent with theory predicting that, under migration–selection balance, adaptive architecture evolves toward tighter linkage rather than remaining polygenic and diffuse. Together, these results indicate that the co-existence of local adaptation in a hierarchically structured, dispersal-prone species depends less on how much dispersal occurs than on whether it results in gene flow, and on the genomic architecture available to resist it once it does. That concentration cuts both ways. An architecture of few, tightly linked loci resist swamping, but it also constrains the response to selection: adaptation from a small number of large-effect regions has less capacity to track a moving optimum than a diffuse polygenic basis (Kardos & Luikart 2021), and the adaptive potential of these populations depends further on their standing variation and effective size, several estimates of which are low here. At a range edge where thermal and hydrological conditions are changing fastest, the question is therefore not only whether local adaptation can survive gene flow, but whether the architecture that allows it to survive also allows it to respond to fast environmental changes.

## Acknowledgements

We thank the “ORE DIAPFC” involved in data collection for the Nivelle/Scorff/Sélune/Bresle rivers, Orekan Gestión Ambiental de Navarra for samples of the Bidasoa, and the fishing societies for the other rivers (especially the Fédérations de pêche du Morbihan, Finistère et Pas de Calais). The samples used in this study were provided by the Biological Resource Centre Colisa (DOI: <u>Biological Resource Centre Colisa</u>) » part of BRC4Env (DOI: https://doi.org/10.15454/TRBJTB), of the Research Infrastructure AgroBRC-RARe.

