## Supplemental Figures and Text for "Dispersal, gene flow and adaptive differentiation in hierarchically structured populations"

### Supplementary Figures

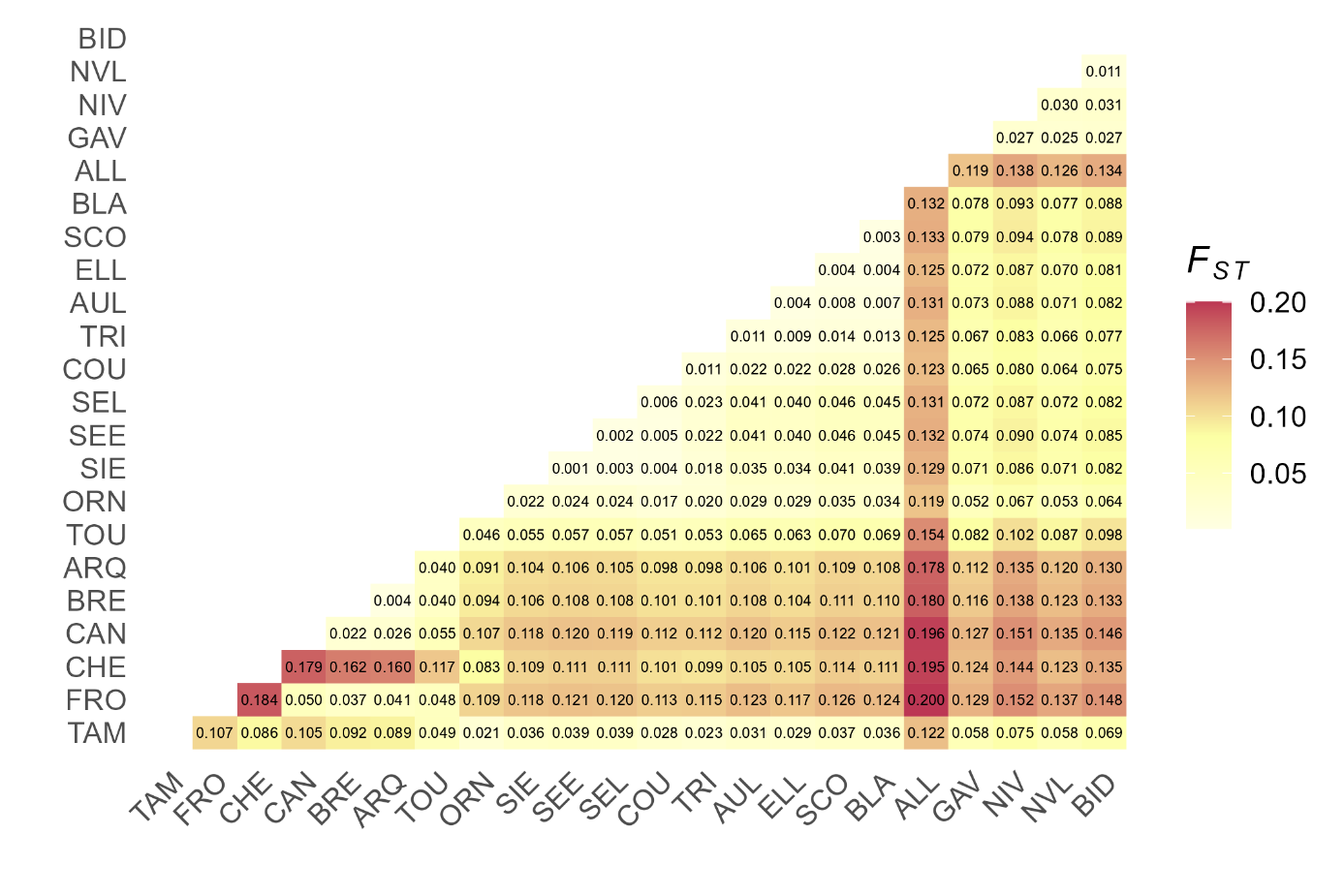

**Fig S1. Pairwise genetic differentiation (*F*ST) matrix estimated using all available SNP markers.** Population codes are shown on the x- and y-axes (see Table 1 for abbreviations), and each cell indicates the *F*_ST_ value calculated between the corresponding pair of populations. The color scale ranges from light yellow (low *F*ST, ≈ 0.00) to dark red (high *F*ST, ≈ 0.20).

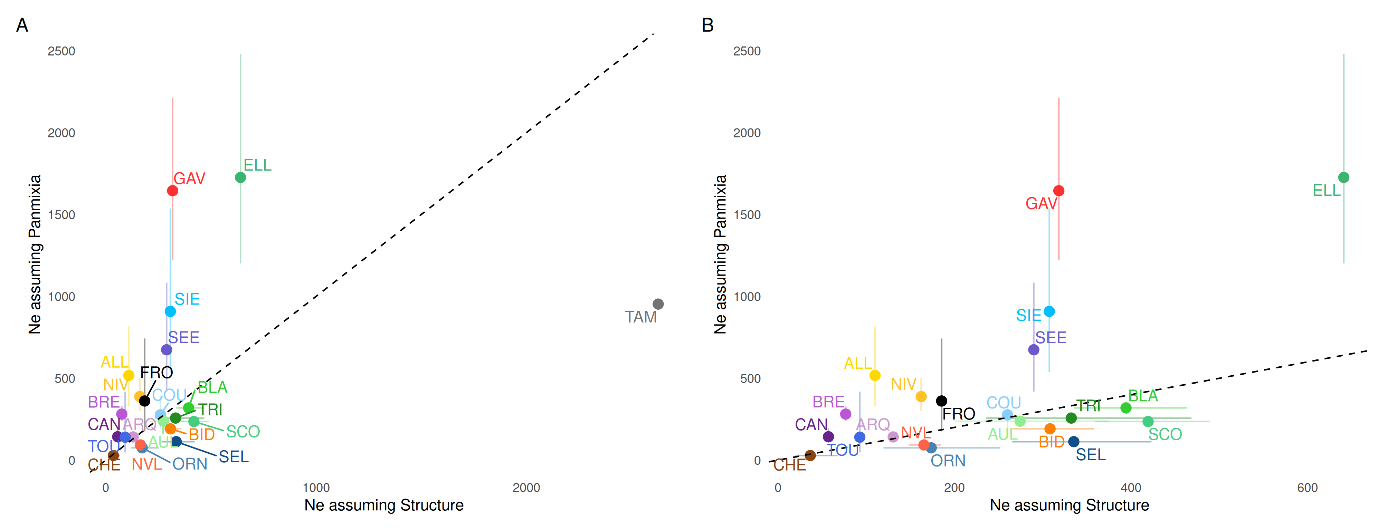

**Fig S2. Comparison of contemporary effective population size (*N*e) estimates obtained assuming population structure (x-axis) versus panmixia (y-axis), based on a linkage-disequilibrium method restricted to loci pairs on different chromosomes to minimize bias from physical linkage.** Points represent individual populations, colour-coded by population acronym; the dashed line indicates the 1:1 identity line. (A) Full dataset including TAM. (B) Populations excluding TAM, shown separately due to its outlying values. Larger *N*e estimates under the structure-assuming option relative to the panmixia estimate (points above the identity line) indicate population subdivision.

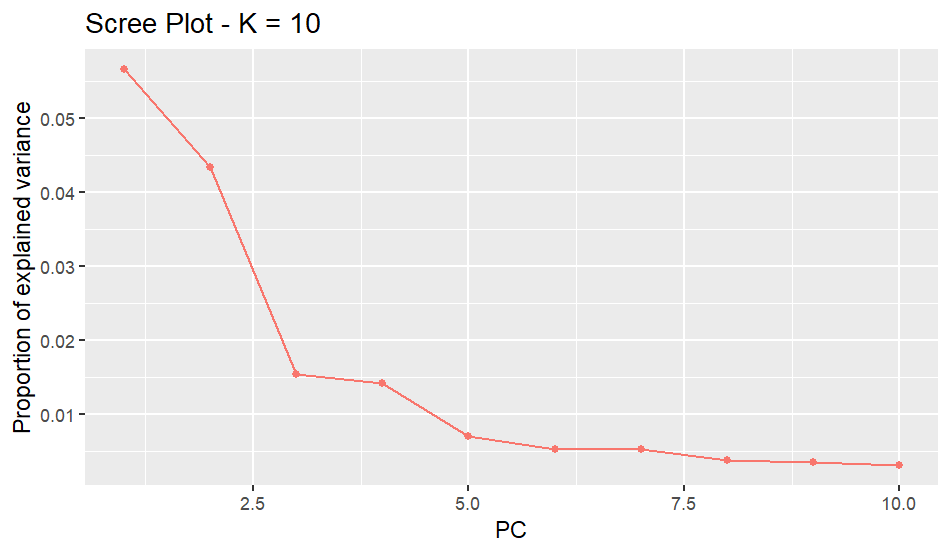

**Fig S3.** **Scree plot of principal components inferred with pcadapt.** Scree plot showing the proportion of genomic variance explained by the first ten principal components computed from the LD‑pruned SNP dataset using pcadapt v.4.4.0. The curve exhibits a steep decline over the first few components followed by a marked inflexion and an approximately linear tail, and application of Cattell’s rule indicated that four PCs (K = 4) captured the major axes of population structure used in subsequent analyses.

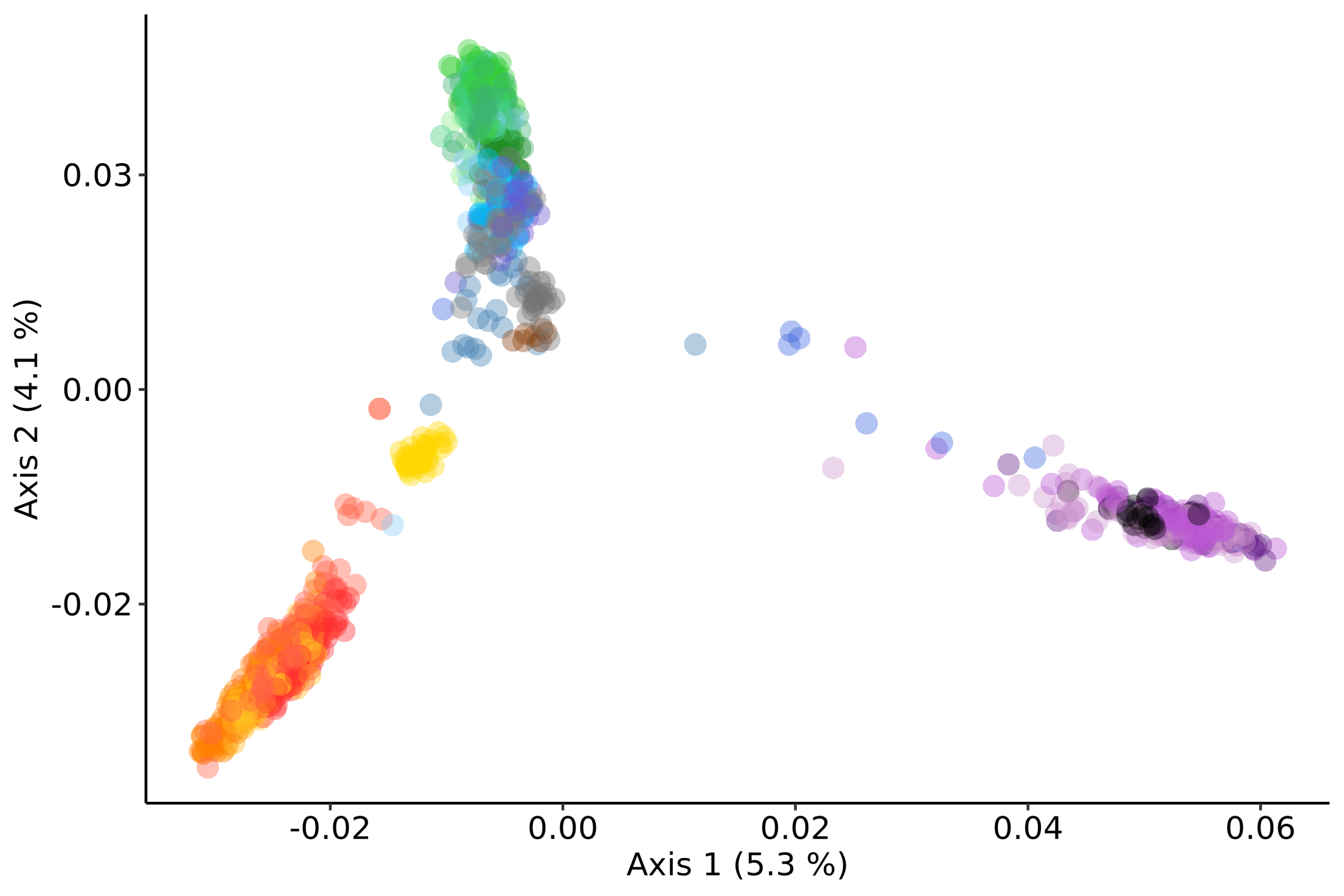

**Fig S4.** **Principal Component Analysis of French Atlantic salmon populations (axes 1 and 2).** Ordination of individual genotypes along PC1 (5.7% of total variance) and PC2 (4.4% of total variance) based on 48,933 LD‑pruned SNPs. Points represent individual Atlantic salmon coloured by river of origin, with population acronyms indicated in the legend (see Table S1 for full names). PC1 primarily separates the English Channel/Upper Normandy and Touques/Frome populations from Brittany, Lower Normandy/Mont Saint Michel Bay and Southwest Atlantic rivers, whereas PC2 further differentiates Brittany and Lower Normandy rivers from the remaining coastal groups.

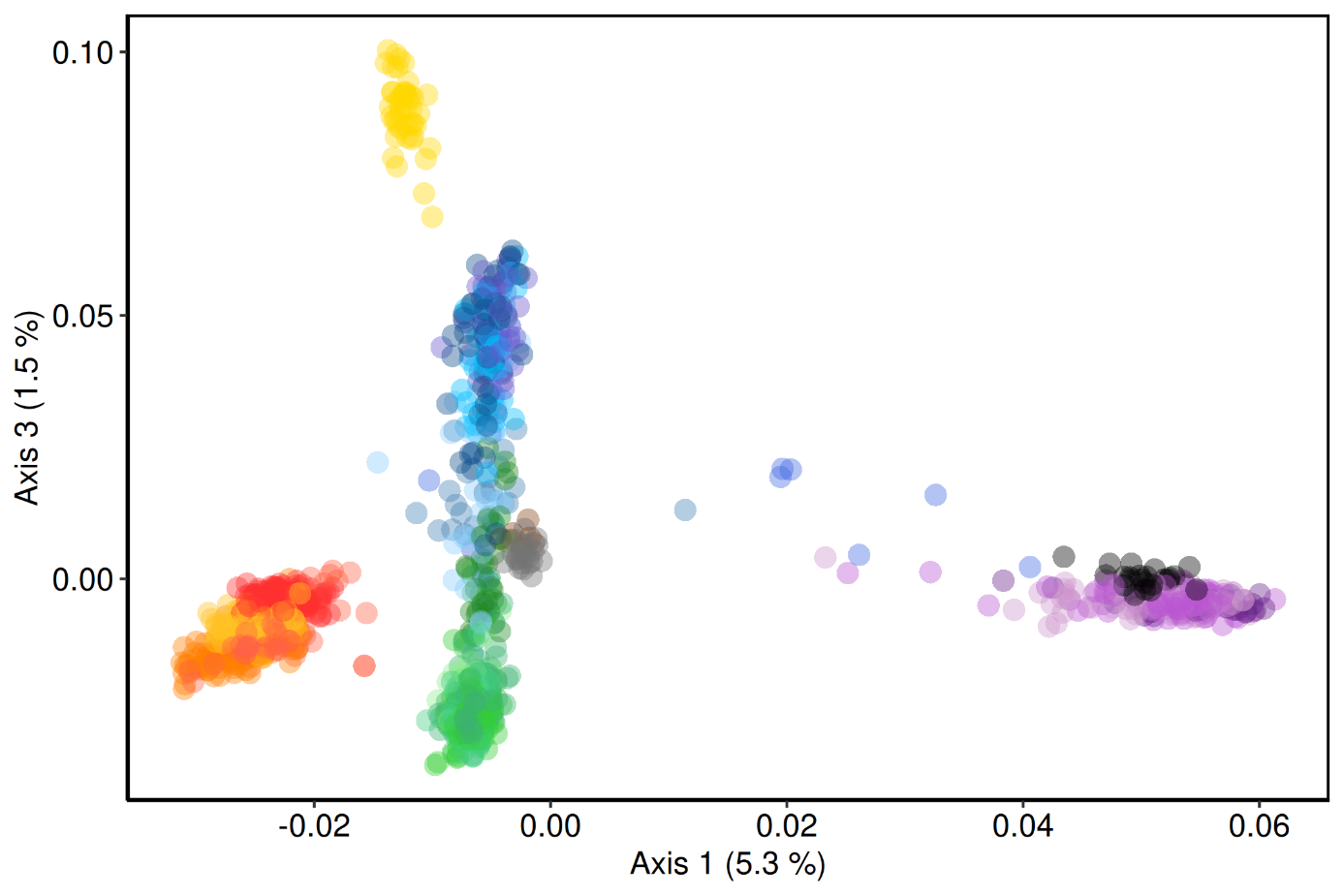

**Fig S5.** **Principal Component Analysis of French Atlantic salmon populations (axes 1 and 3).** Ordination of individual genotypes along PC1 (5.3–5.7% of total variance; same axis as in Fig. S4) and PC3 (1.5% of total variance) based on 48,933 LD‑pruned SNPs. Points represent individual Atlantic salmon coloured by river of origin, as in Fig. S4. PC3 highlights the strong divergence of the Allier population (ALL) and increases separation between Southwest Atlantic rivers and the English Channel/Upper Normandy group, while individuals from Brittany and Lower Normandy occupy an intermediate position along PC3.

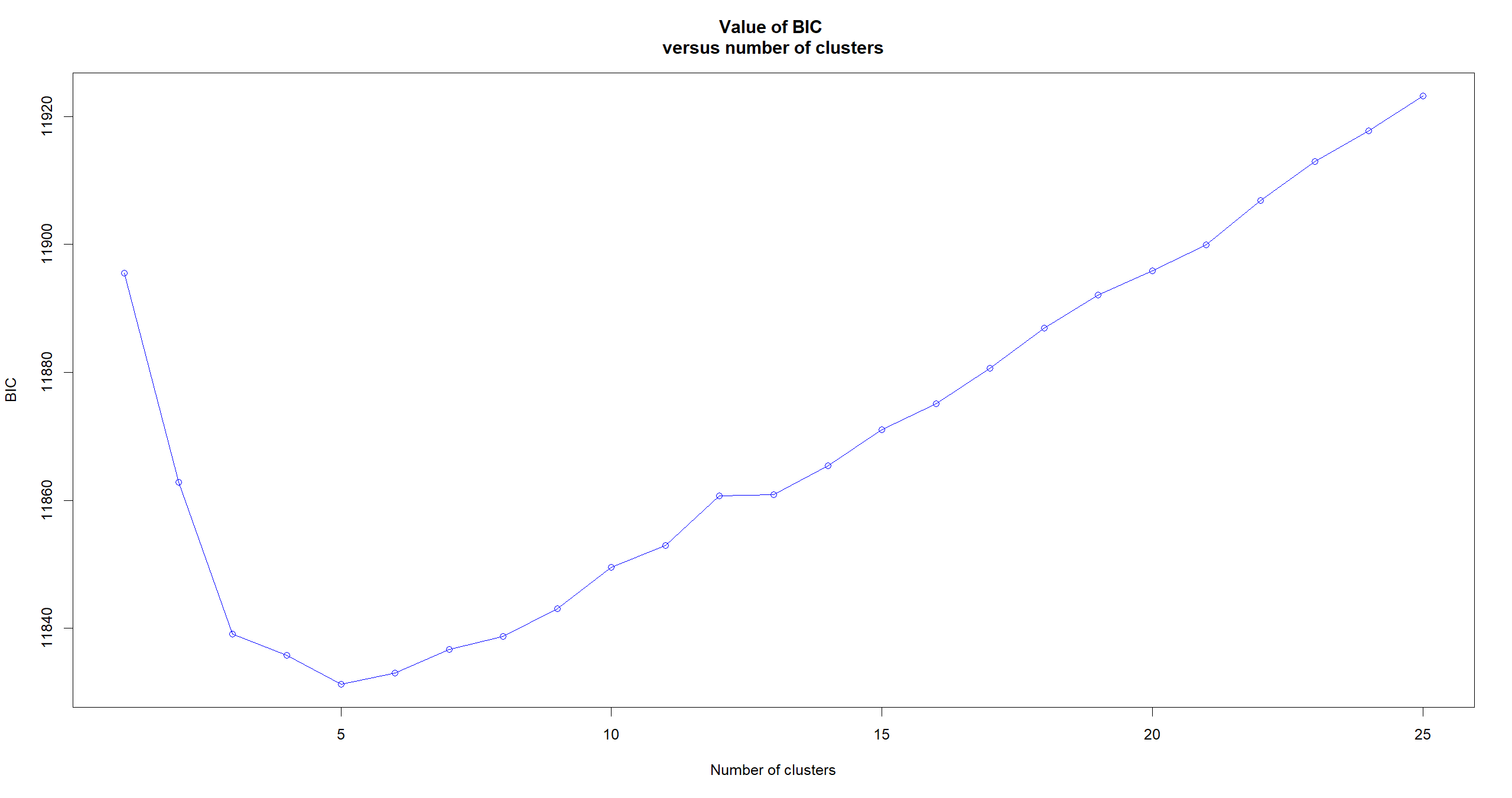

**Fig. S6. Bayesian Information Criterion (BIC) for DAPC cluster optimisation.** Plot of BIC values as a function of the number of clusters (K) obtained with the find.clusters function in adegenet, based on successive K‑means clustering of the LD‑pruned SNP dataset after PCA transformation. The BIC decreases sharply from low K and reaches a clear minimum around five clusters, after which additional clusters only worsen the model fit, supporting the retention of K = 5 as the optimal number of genetic clusters used in subsequent DAPC analyses.

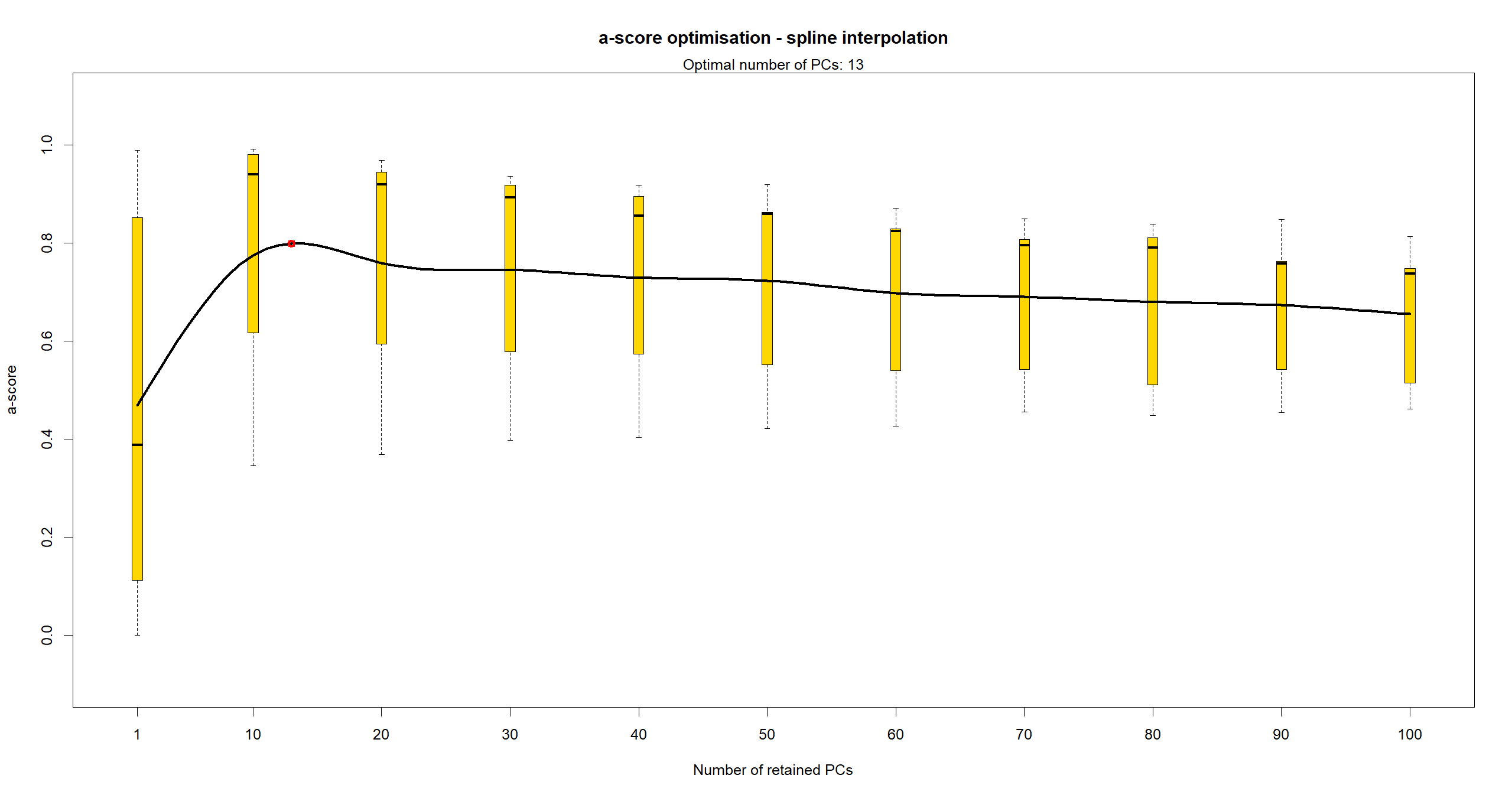

**Fig. S7. α‑score optimisation for the number of principal components retained in DAPC.** Results of the α‑score optimisation procedure implemented in adegenet for the Discriminant Analysis of Principal Components. Boxplots show the distribution of α‑scores (discriminant power corrected for overfitting) for increasing numbers of retained PCs, and the spline‑interpolated curve highlights the trade‑off between discrimination and overfitting. The α‑score reaches a maximum around 13 retained PCs (red point), which was therefore chosen for the final DAPC, ensuring robust discrimination of genetic clusters without inflating spurious structure.

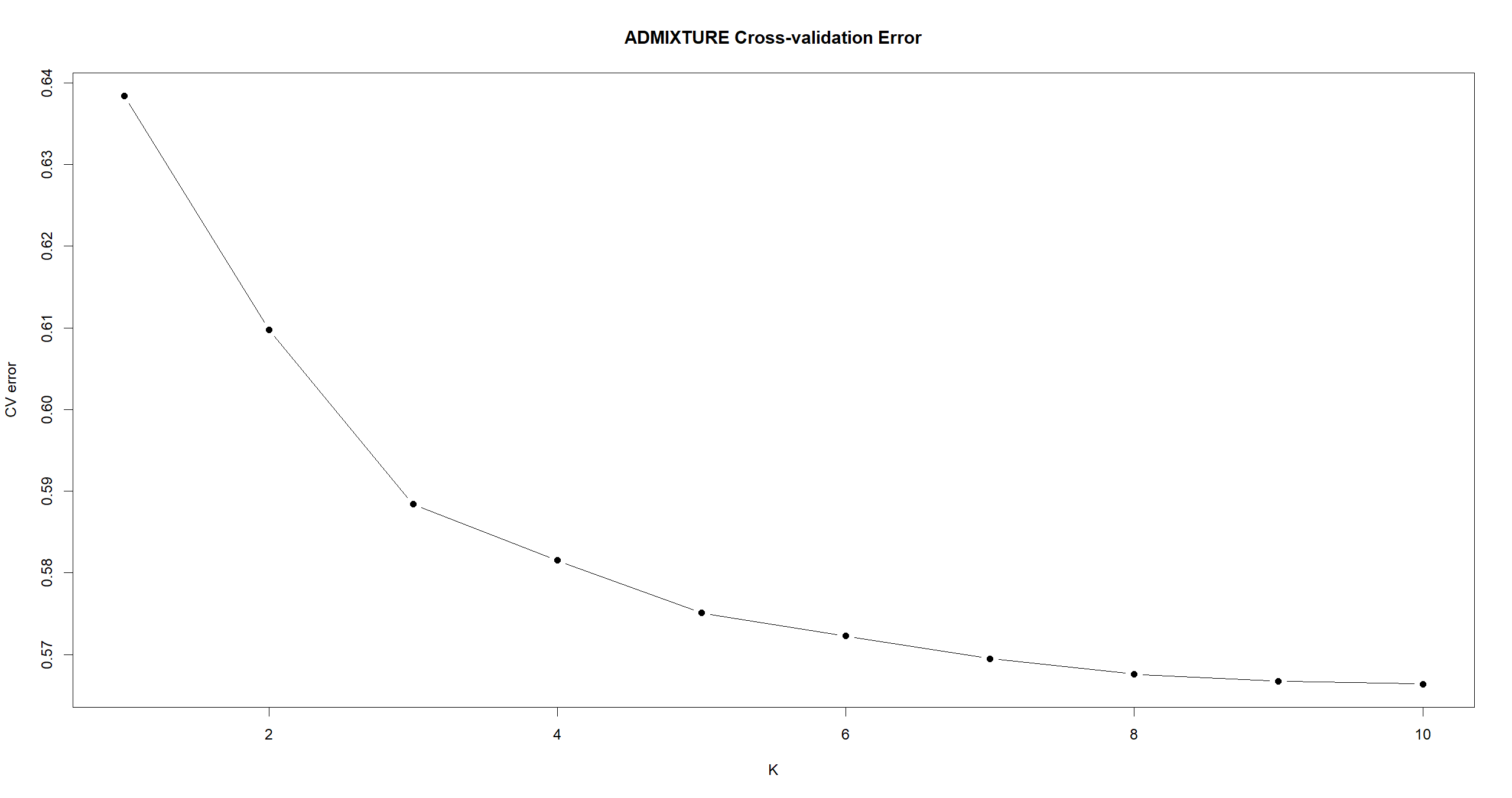

**Fig. S8. ADMIXTURE cross‑validation error as a function of K.** Cross‑validation errors obtained from unsupervised ADMIXTURE runs for K = 1–10 ancestral populations using the --cv option. Each point represents the mean cross‑validation error for a given K, and the curve shows a pronounced decline between K = 1 and K = 5 followed by a shallow plateau, indicating that models with five clusters provide the best compromise between goodness‑of‑fit and parsimony and supporting K = 5 as the most likely number of main genetic clusters.

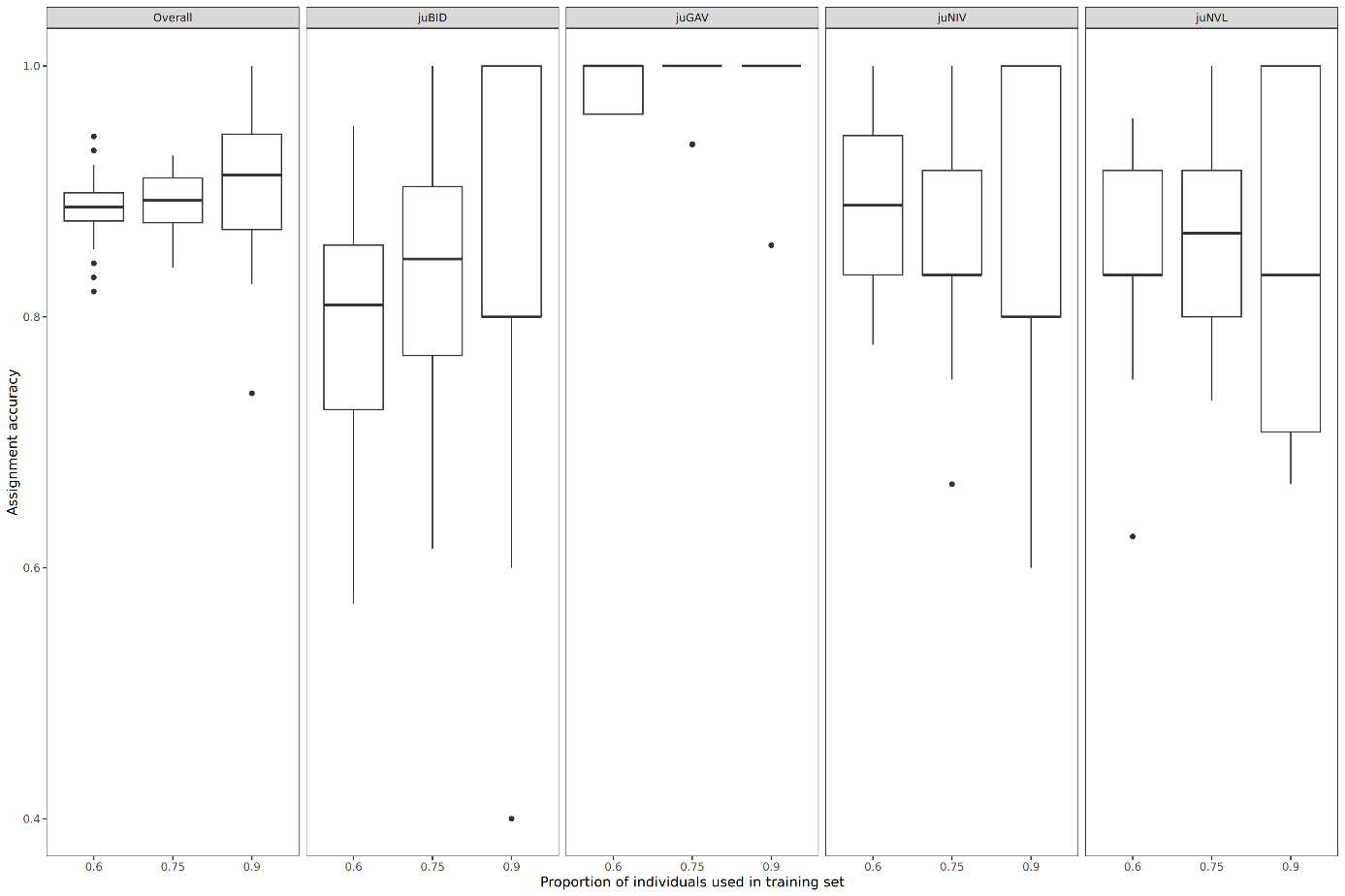

**Fig. S9.** **Assignment accuracies estimated via Monte-Carlo cross-validation with a Support Vector Machine (svm) classification method for the Adour regional group.** We used three levels of training sets (50%, 70% and 90% of individuals from each population, on x-axis). ‘Overall’ represents the whole dataset, composed of 225 individuals (53, 65, 46 and 61 individuals from left to right). See Table 1 for abbreviations. We used all available markers (48933 SNPs) for each combination of training sets.

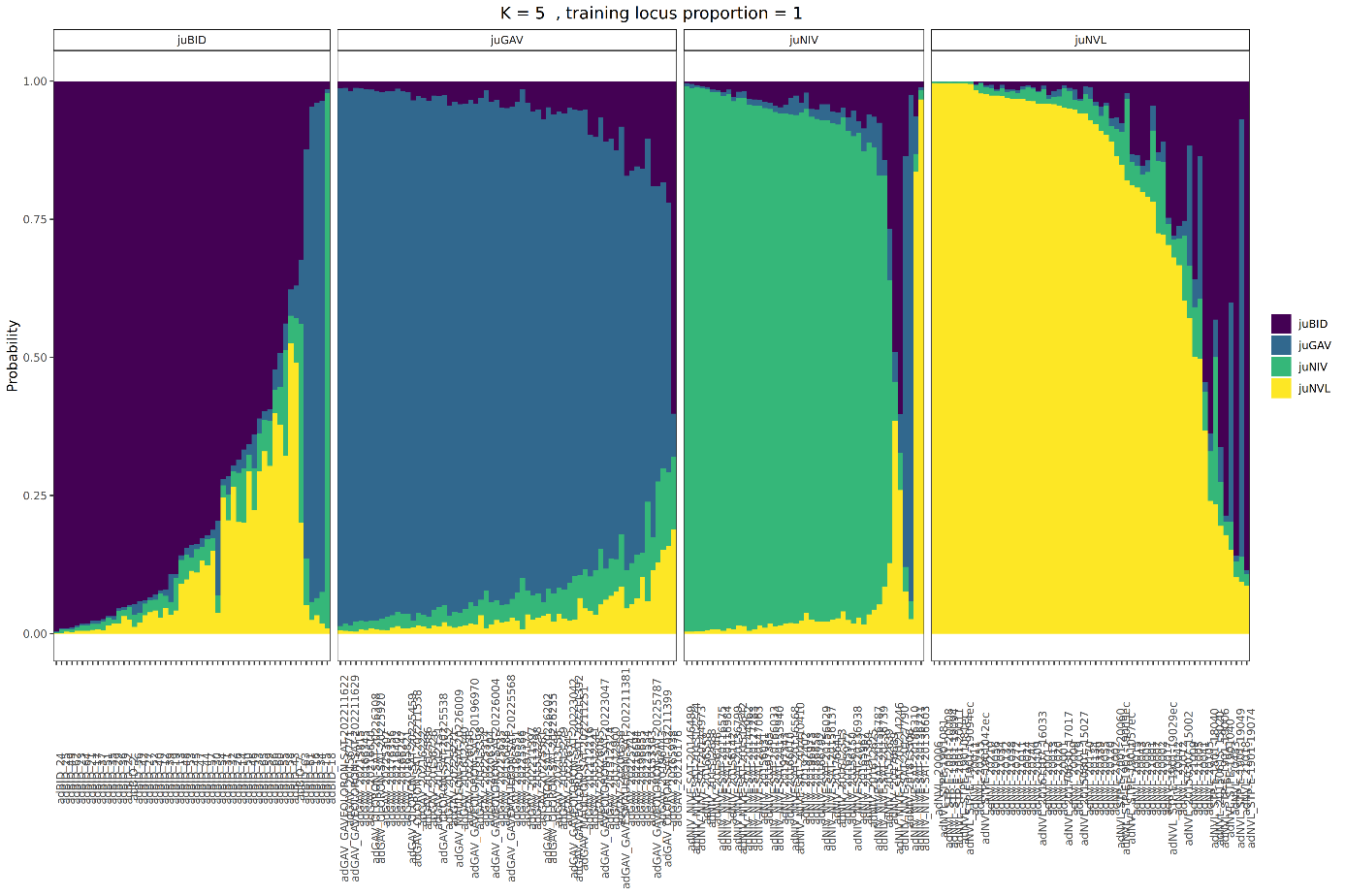

**Fig. S10.** **Membership probabilities estimated via K-fold cross-validation with a Support Vector Machine (svm) classification method for the Adour regional group.** We estimated the results via 5-fold cross validation using all available markers. Each panel, from left to right, was composed of 53, 65, 46 and 61 individuals respectively.

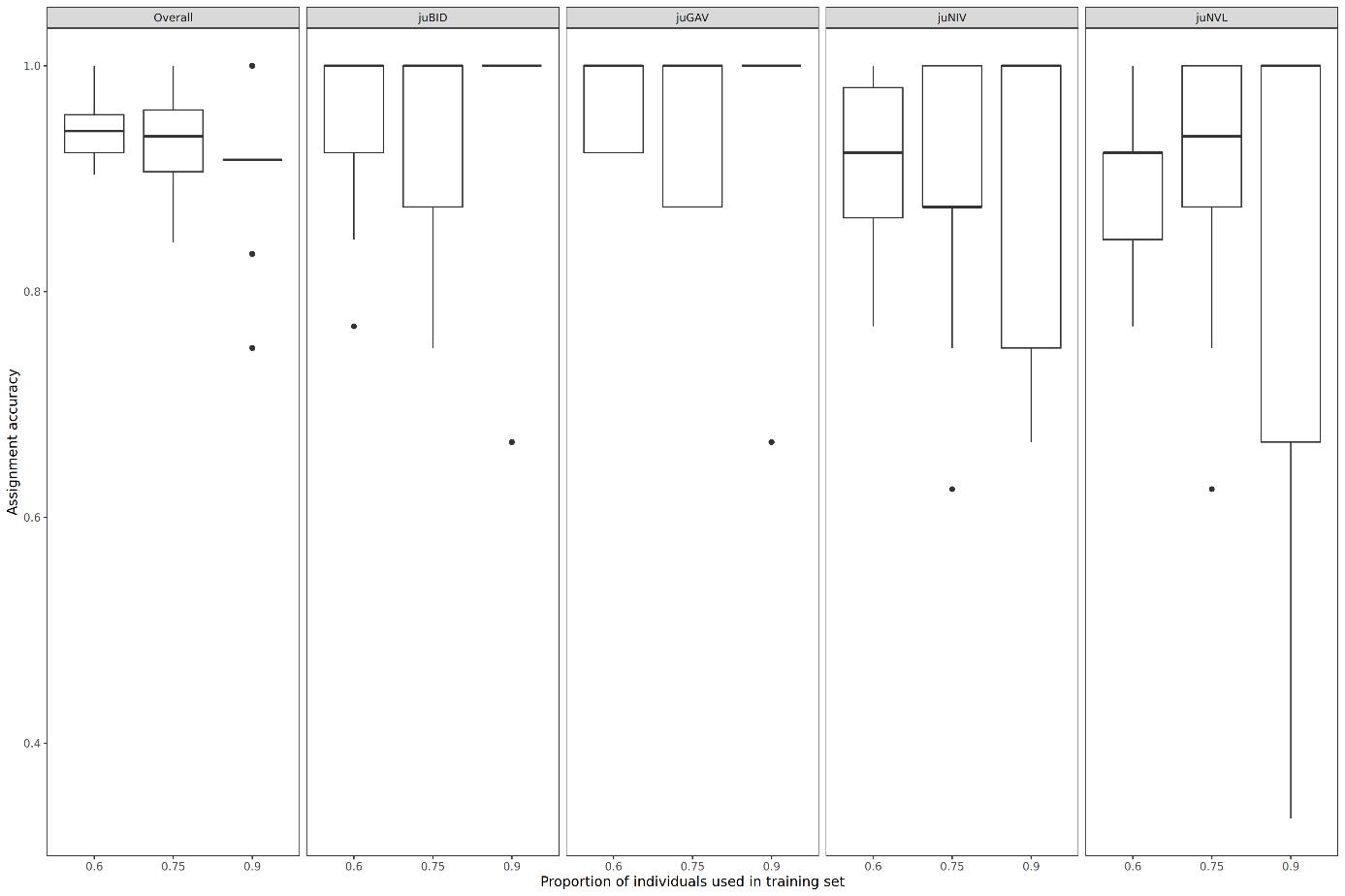

**Fig. S11.** **Assignment accuracies estimated via Monte-Carlo cross-validation with a Support Vector Machine (svm) classification method for the Adour regional group after training the baseline data.** We used three levels of training sets (50%, 70% and 90% of individuals from each population, on x-axis). ‘Overall’ represents the whole dataset, composed of 132 individuals (33 for each panel). See Table 1 for abbreviations. We used all available markers (48933 SNPs) for each combination of training sets.

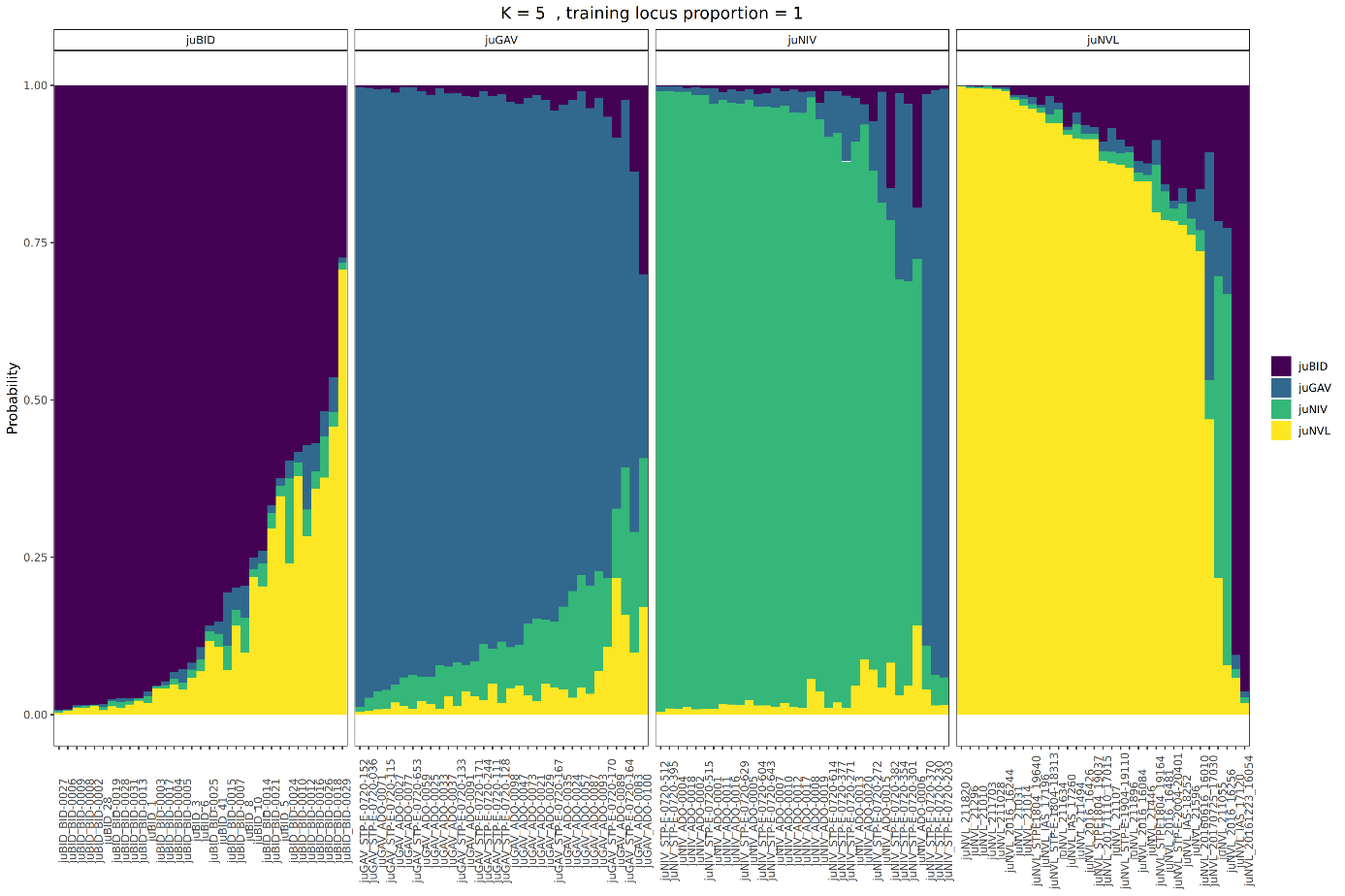

**Fig. S12.** **Membership probabilities estimated via K-fold cross-validation with a Support Vector Machine (svm) classification method for the Adour regional group after training the baseline data.** We estimated the results via 5-fold cross validation using all available markers. Each panel, from left to right, was composed of 33 individuals respectively.

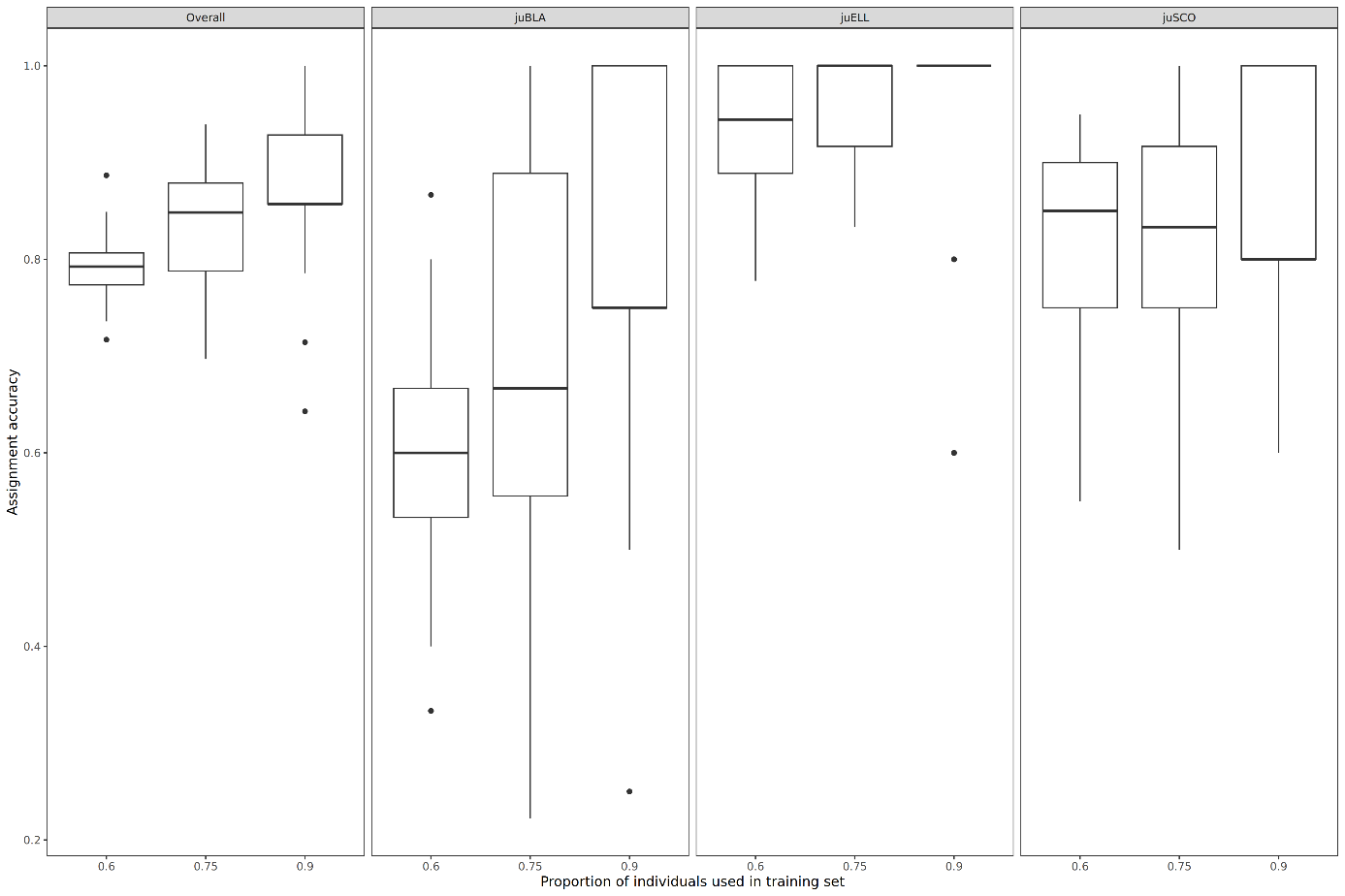

**Fig. S13.** **Assignment accuracies estimated via Monte-Carlo cross-validation with a Support Vector Machine (svm) classification method for the Brittany regional group.** We used three levels of training sets (50%, 70% and 90% of individuals from each population, on x-axis). ‘Overall’ represents the whole dataset, composed of 133 individuals (37, 46 and 50 individuals from left to right). See Table 1 for abbreviations. We used all available markers (48933 SNPs) for each combination of training sets.

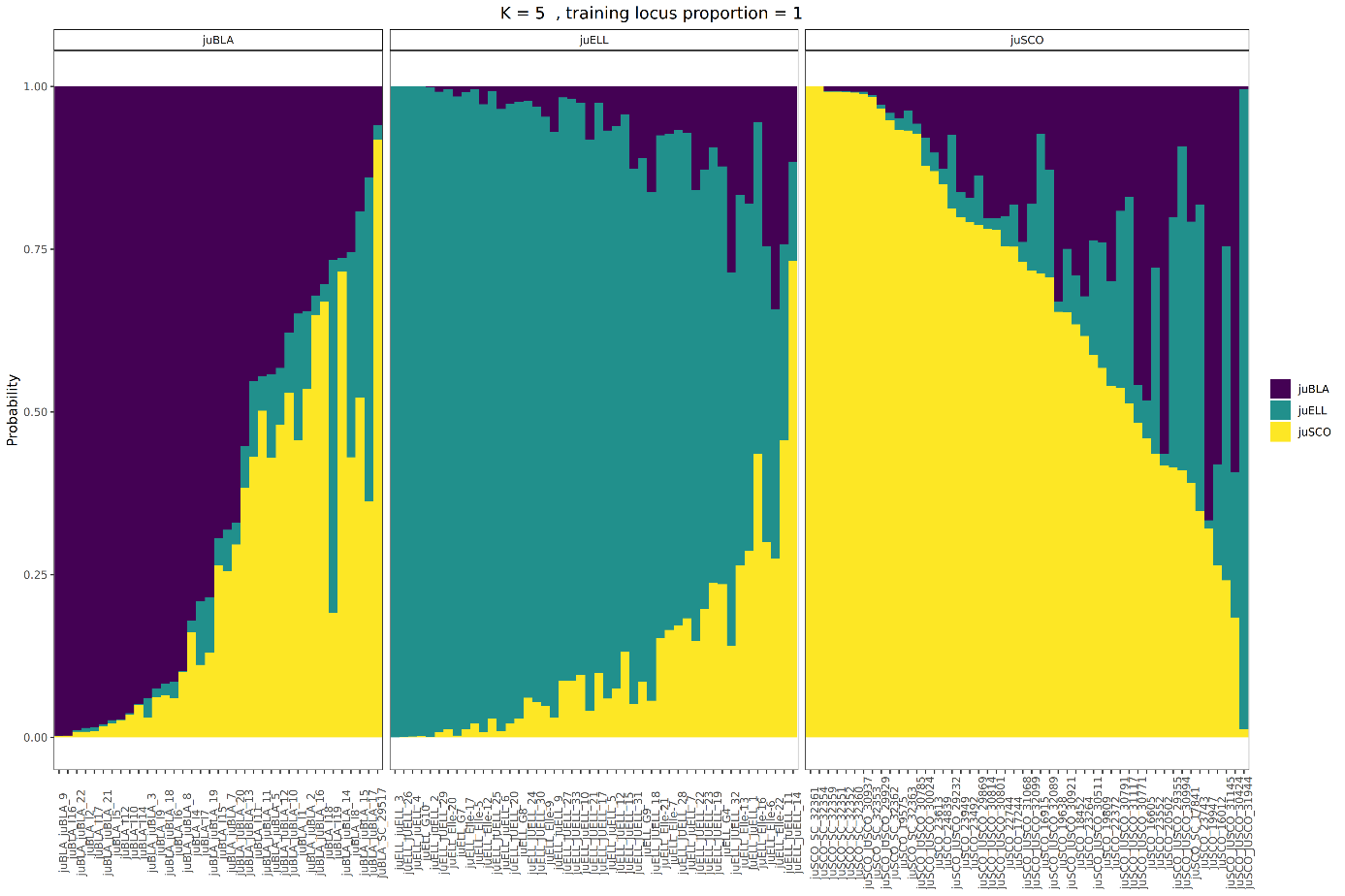

**Fig. S14.** **Membership probabilities estimated via K-fold cross-validation with a Support Vector Machine (svm) classification method for the Brittany regional group.** We estimated the results via 5-fold cross validation using all available markers. Each panel, from left to right, was composed of 37, 46 and 50 individuals respectively.

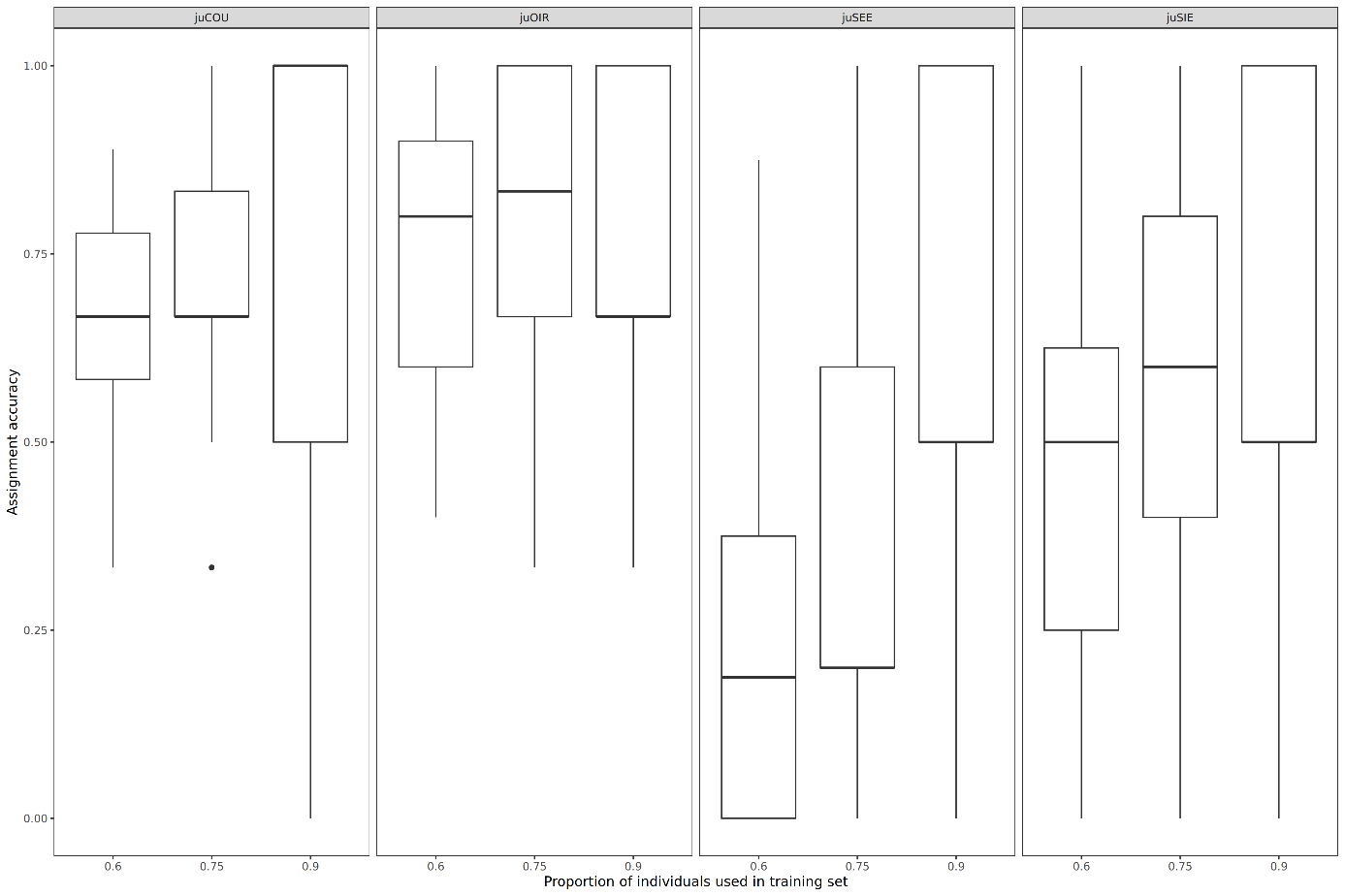

**Fig. S15.** **Assignment accuracies estimated via Monte-Carlo cross-validation with a Support Vector Machine (svm) classification method for the Lower Normandy regional group.** We used three levels of training sets (50%, 70% and 90% of individuals from each population, on x-axis). ‘Overall’ represents the whole dataset, composed of 88 individuals (23, 26, 19 and 20 individuals from left to right). See Table 1 for abbreviations. We used all available markers (48933 SNPs) for each combination of training sets.

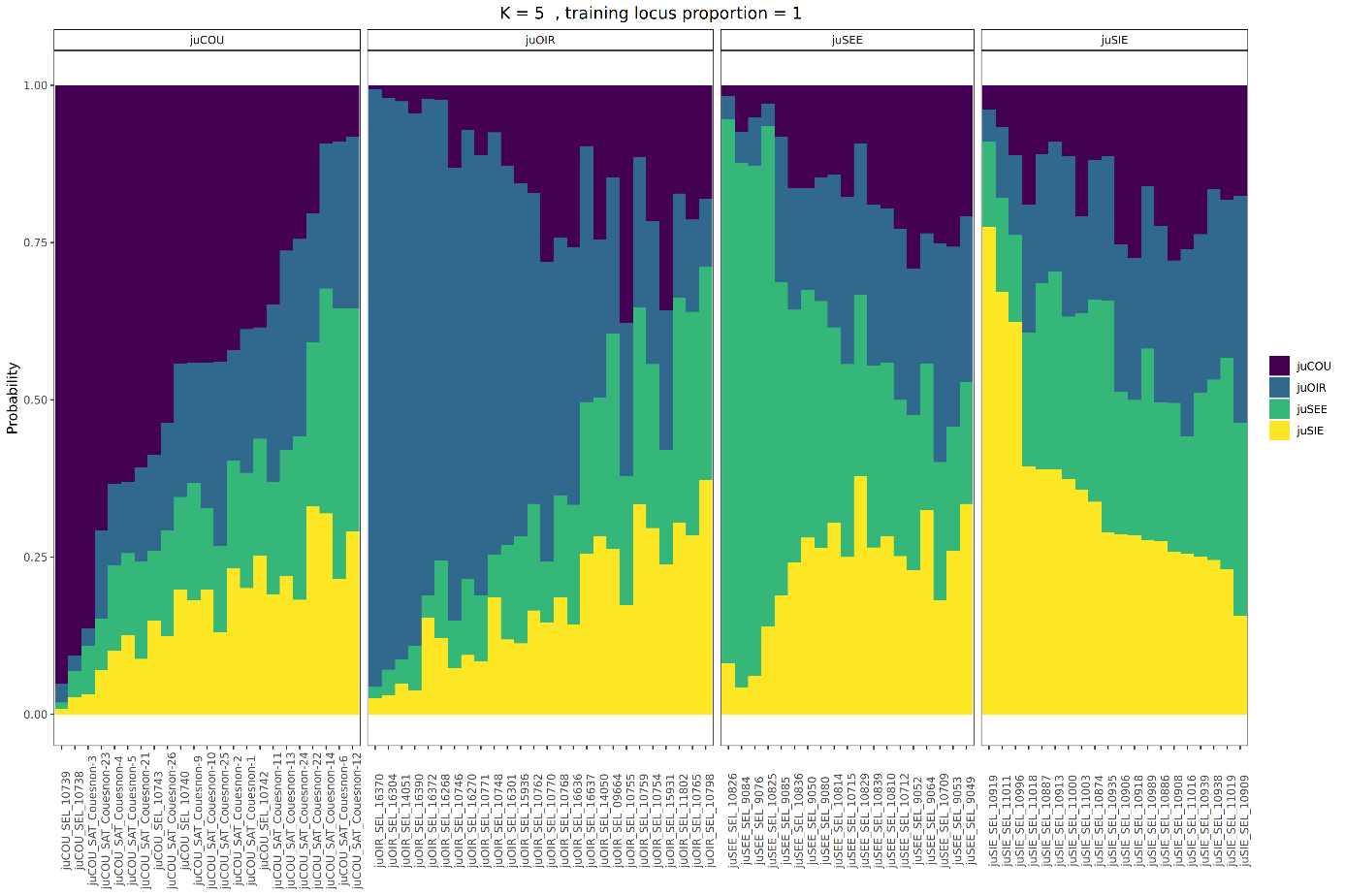

**Fig. S16.** **Membership probabilities estimated via K-fold cross-validation with a Support Vector Machine (svm) classification method for the Lower Normandy regional group.** We estimated the results via 5-fold cross validation using all available markers. Each panel, from left to right, was composed of 23, 26, 19 and 20 individuals respectively.

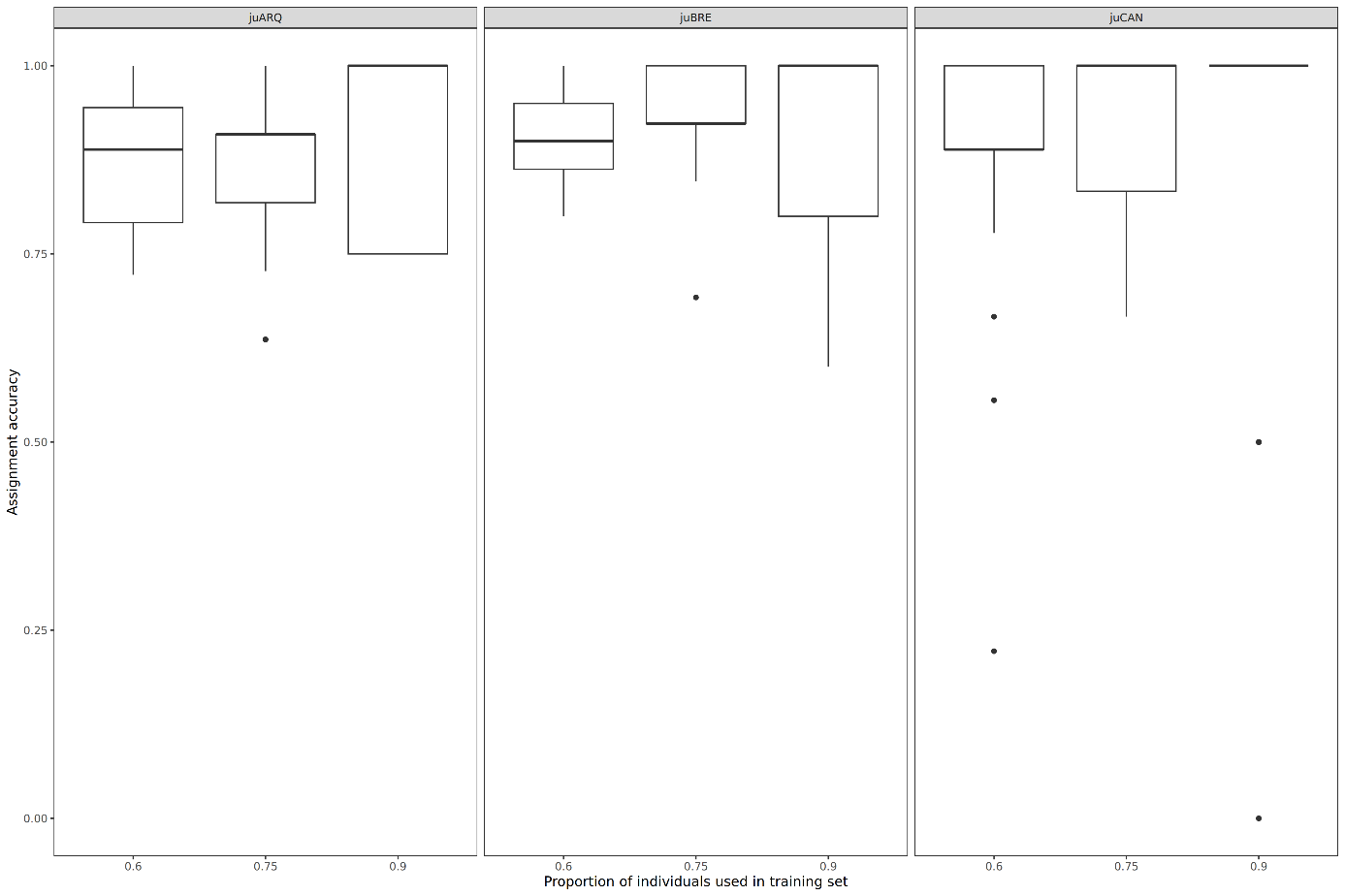

**Fig. S17.** **Assignment accuracies estimated via Monte-Carlo cross-validation with a Support Vector Machine (svm) classification method for the Upper Normandy regional group.** We used three levels of training sets (50%, 70% and 90% of individuals from each population, on x-axis). ‘Overall’ represents the whole dataset, composed of 117 individuals (44, 51 and 22 individuals from left to right). See Table 1 for abbreviations. We used all available markers (48933 SNPs) for each combination of training sets.

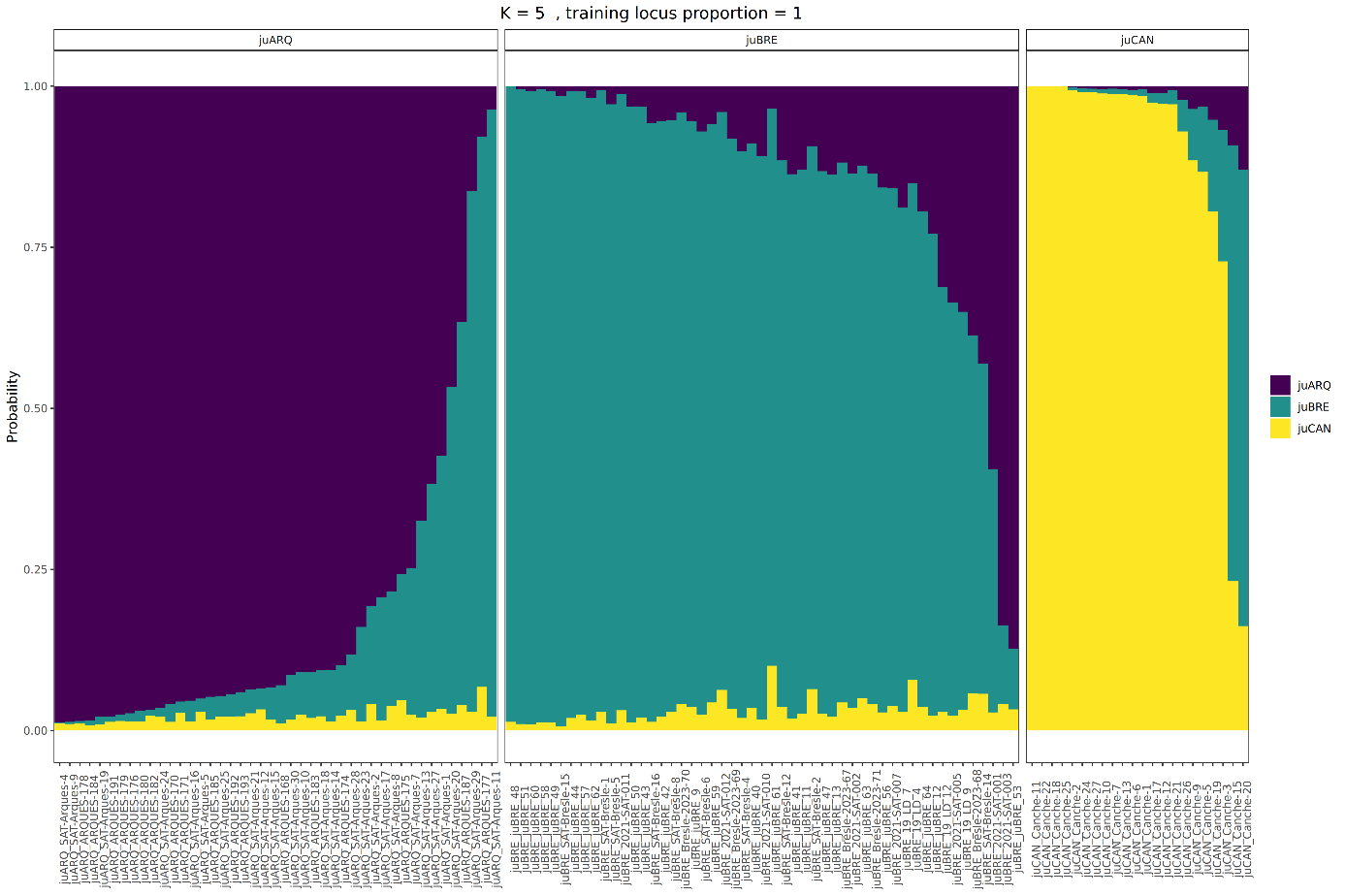

**Fig. S18.** **Membership probabilities estimated via K-fold cross-validation with a Support Vector Machine (svm) classification method for the Upper Normandy regional group.** We estimated the results via 5-fold cross validation using all available markers. Each panel, from left to right, was composed of 44, 51 and 22 individuals respectively.

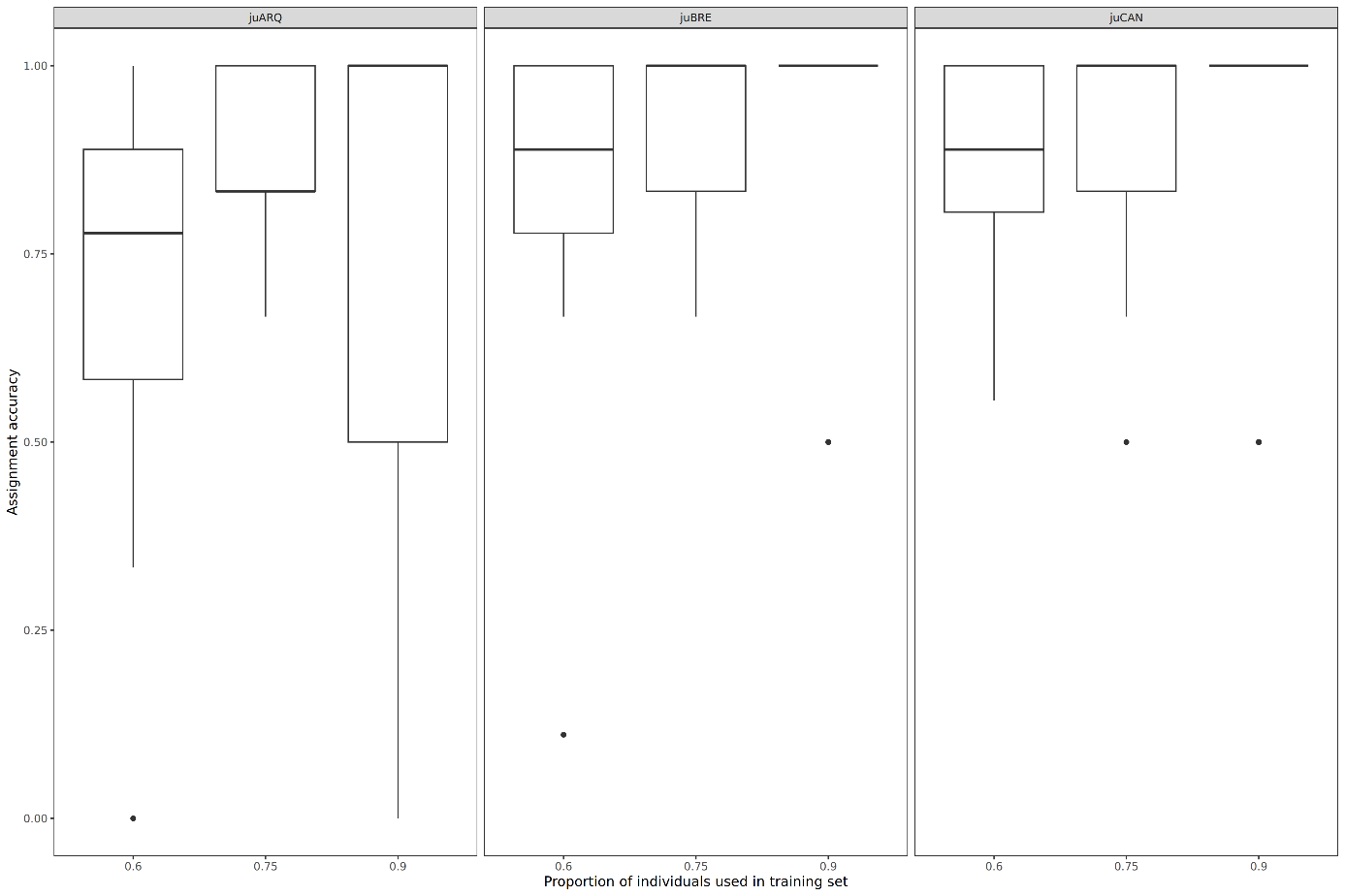

**Fig. S19.** **Assignment accuracies estimated via Monte-Carlo cross-validation with a Support Vector Machine (svm) classification method for the Upper Normandy regional group after training the baseline data.** We used three levels of training sets (50%, 70% and 90% of individuals from each population, on x-axis). ‘Overall’ represents the whole dataset, composed of 66 individuals (22 for each panel). See Table 1 for abbreviations. We used all available markers (48933 SNPs) for each combination of training sets.

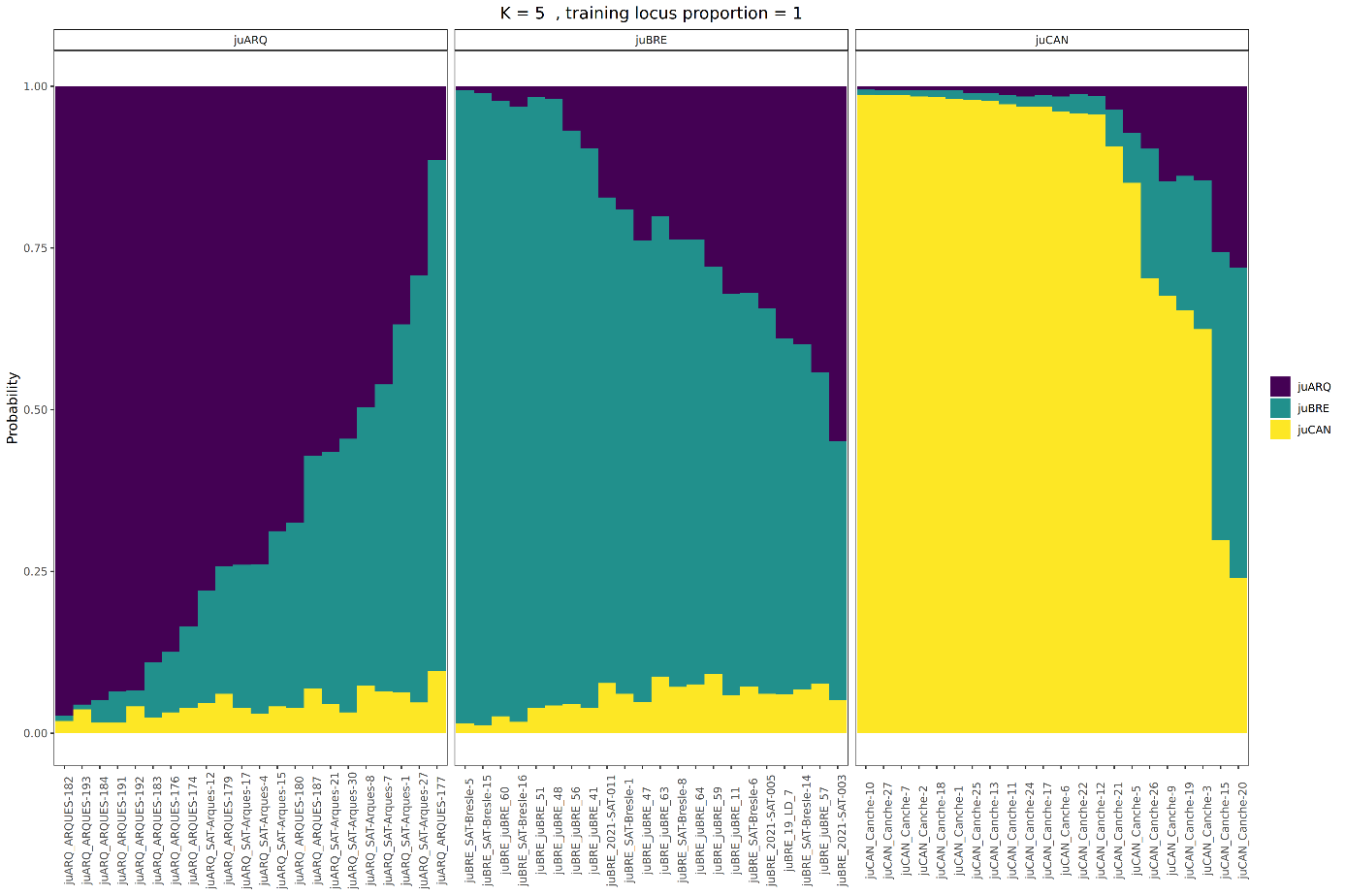

**Fig. S20.** **Membership probabilities estimated via K-fold cross-validation with a Support Vector Machine (svm) classification method for the Upper Normandy regional group after training the baseline data.** We estimated the results via 5-fold cross validation using all available markers. Each panel, from left to right, was composed of 22 individuals respectively.

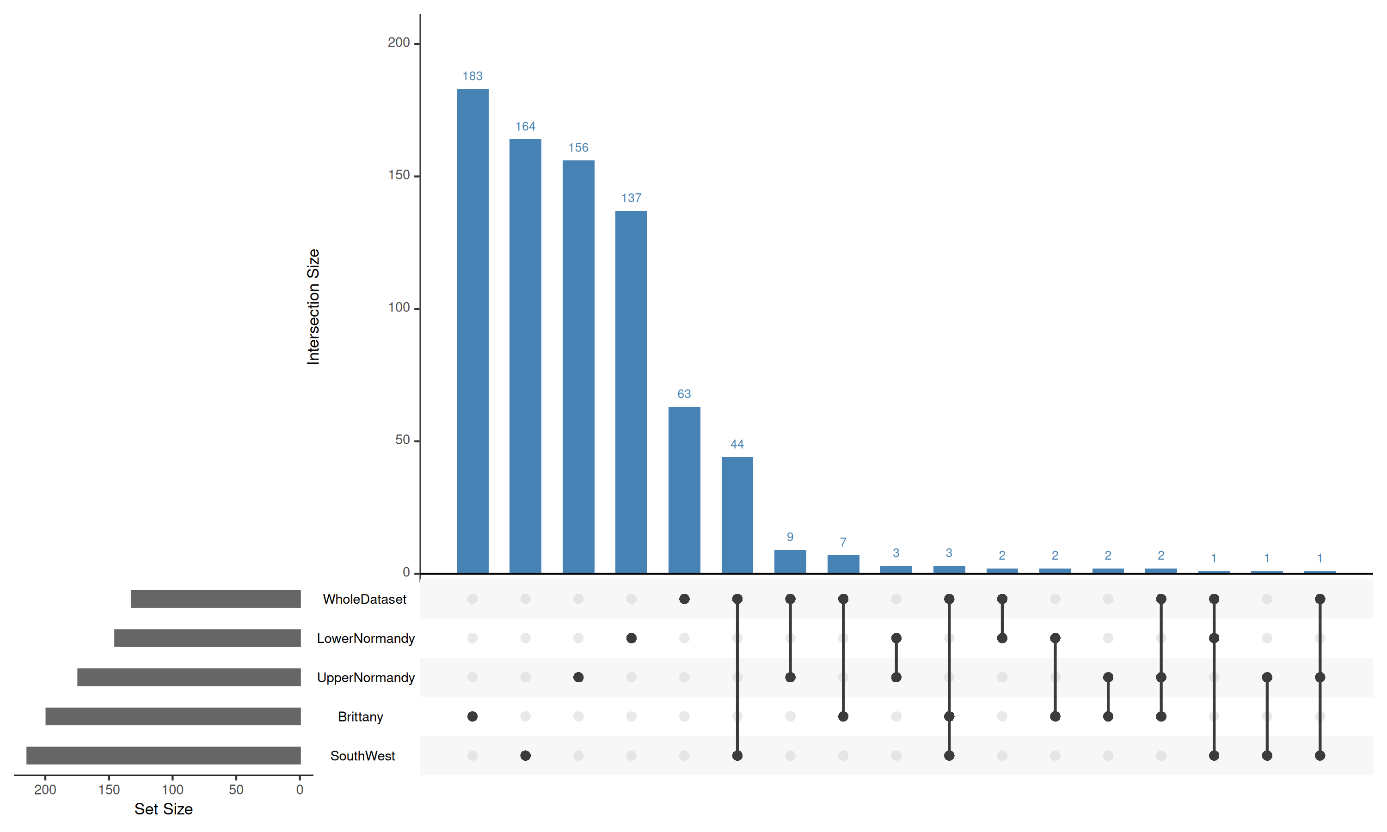

**Fig. S21. UpSet plot of XtX outlier SNP overlap between the national-scale and metapopulation-specific genome scans.** Horizontal bars (bottom left) show the total number of outlier SNPs identified in each of the five XtX scans: Southwest (n = 214), Brittany (n = 199), Upper Normandy (n = 174), Lower Normandy (n = 145), and the national-scale whole-dataset scan (WholeDataset, n = 132). Vertical bars (top) show the number of SNPs in each intersection, defined by the connected dots in the matrix below; single dots indicate SNPs detected exclusively in that scan, whereas connected dots indicate SNPs shared between the corresponding scans. Intersections are ordered by decreasing size.

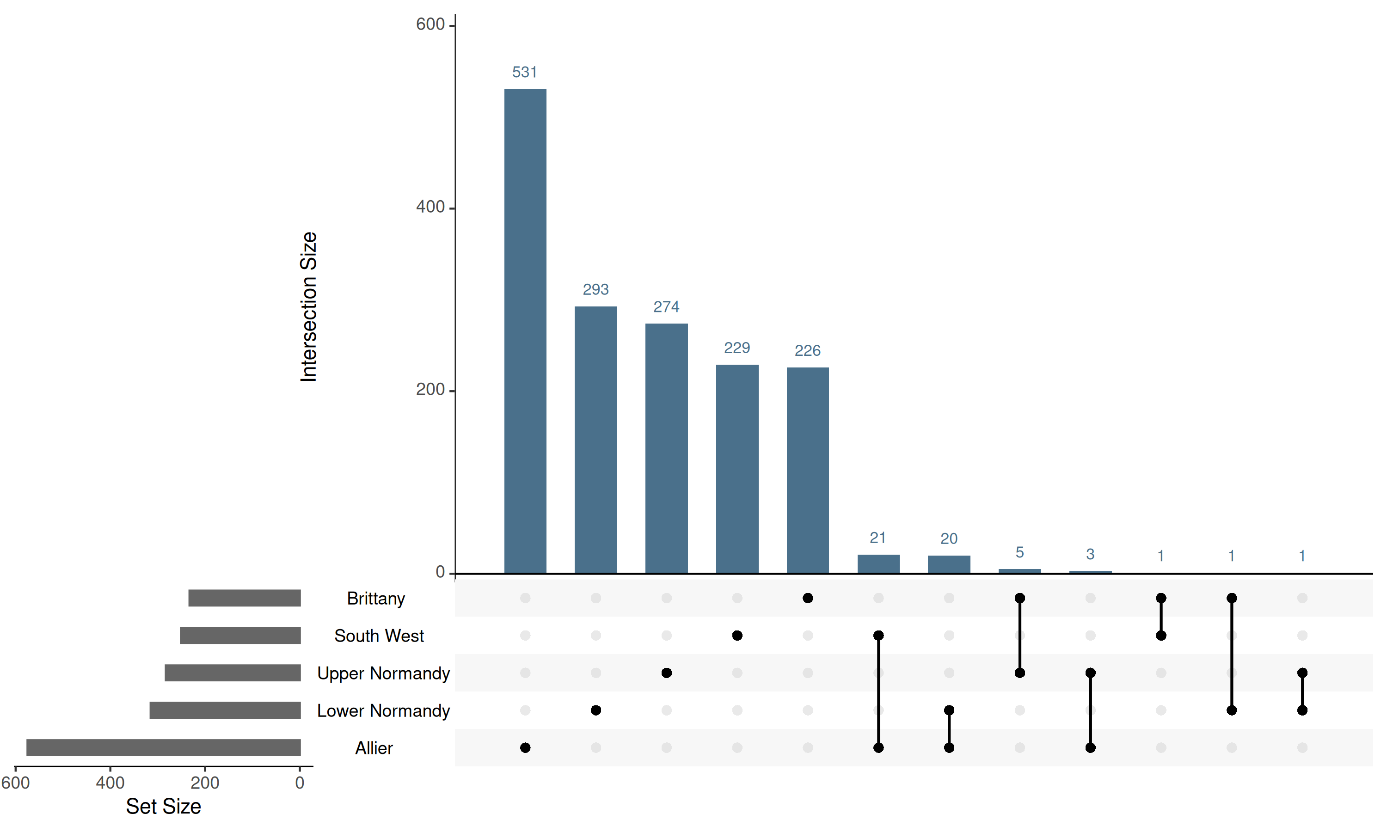

**Fig. S22. UpSet plot of C2 outlier SNP overlap between the five regional one-vs-rest contrasts.** Horizontal bars (bottom left) show the total number of outlier SNPs identified in each C2 contrast: Allier (n = 575), Lower Normandy (n = 315), Upper Normandy (n = 283), Southwest (n = 251), and Brittany (n = 233). Vertical bars (top) show the number of SNPs in each intersection, defined by the connected dots in the matrix below; single dots indicate SNPs detected exclusively in that contrast, whereas connected dots indicate SNPs shared between the corresponding contrasts. Intersections are ordered by decreasing size.

### Supplementary Text

#### Extraction of climatic and physiographic variables

**Temperature data acquisition and processing**

Unlike studies relying on global gridded climate datasets (e.g., WorldClim bioclimatic variables) as proxies for river thermal conditions, our approach is based on directly measured water temperatures, which more accurately reflect the thermal environment experienced by Atlantic salmon populations. We compiled mean water temperatures for all studied rivers over a 15-year period spanning January 2011 to December 2025, providing a robust interannual average representative of contemporary thermal conditions at each site. For the majority of rivers, temperature records were automatically extracted from the French national water quality database Naïades (naiades.eaufrance.fr) using the R package *hubeau*. When temperature stations were unavailable on the main river stem, we utilised data from well-monitored tributaries or geographically adjacent proxy rivers (e.g., the Ternoise for the Canche, and the Sélune for the Oir).

To account for sensor measurement errors and technical anomalies (e.g., non-physiological values such as −40°C), we applied a strict quality control filter to the raw data, retaining only temperature readings within a realistic biological and physical range (0–30°C) before computing the interannual mean for each river.

For rivers lacking adequate Naïades coverage, we integrated alternative verified sources. Temperature datasets for the Bresle and Nivelle rivers were obtained from the ORE DIAPFC observatory (https://diapfc.hub.inrae.fr/) and averaged over the same 2011–2025 timeframe. For the Bidasoa River, we used the mean annual water temperature (15.2°C) reported by García-Vega et al. (2025), who sourced this value from the Government of Navarre (2022).

**River length and geographic distance**

We obtained river length data primarily from Perrier et al. (2011), and from (García‐Vega *et al.* 2025) for the Bidasoa River. To calculate geographic distance, we measured the distance from the northernmost river (the Canche), following the methodology of Perrier et al. (2011). For the Nivelle and Bidasoa rivers, we manually measured the inter-river distance

**Geological and environmental variables**

We extracted several geological variables using the GeoFRESH platform (Domisch et al., 2024), which integrates, processes, manages, and visualizes standardized spatiotemporal freshwater-related Earth system data. From the soil dataset provided by Hengl et al. (2017), we extracted three specific variables for 2016: clay content (*clyppt*), soil pH ×10 (*phihox*), and coarse fragment volumetric content (*crfvol*). Finally, we extracted mean elevation (*elev*) from the hydrographic dataset developed by Yamazaki et al. (2017).

The selected geological variables were extracted from the upstream catchment of each sampling site (mean of sub-catchment means).
